# Comparative modes of chromatin engagement by PAX::FOXO1 fusions in rhabdomyosarcoma

**DOI:** 10.64898/2026.02.18.706582

**Authors:** Alexi Tallan, Jack Kucinski, Andrew M. Vontell, Chamithi Karunanayake, Rachel A. Hoffman, Benjamin D. Sunkel, Cenny Taslim, Genevieve C. Kendall, Benjamin Z. Stanton

**Affiliations:** Nationwide Children’s Hospital, Center for Childhood Cancer Research, Columbus, OH, USA., 20892, USA; Molecular, Cellular, and Developmental Biology Program, The Ohio State University, Columbus, OH 43210, USA; Department of Pediatrics, The Ohio State University College of Medicine, Columbus, OH, USA; Department of Biological Chemistry & Pharmacology, The Ohio State University College of Medicine, Columbus, OH, USA

**Keywords:** Rare disease epigenetics, Fusion oncoproteins, Childhood cancer etiology, Epigenetic mechanisms, Cancer epigenomics, Pioneer function, Sarcoma

## Abstract

Fusion positive rhabdomyosarcoma (FP-RMS) is an aggressive soft-tissue sarcoma that most frequently affects children and adolescents. Treatment options and outcomes for children with this cancer remain poor, non-specific, and broadly toxic despite decades of research. The defining molecular drivers of the more aggressive fusion-positive subtype of the disease arise from chromosomal translocations that fuse PAX3 or PAX7 to FOXO1 to form PAX3::FOXO1 or PAX7::FOXO1, encoding fusion oncoprotein transcription factors. Despite their high degree of similarity, *PAX3::FOXO1* correlates with worse patient overall survival than *PAX7::FOXO1*. Previous work from our groups and others has revealed evidence focused in chromatin accessibility contexts that PAX3::FOXO1 has key characteristics of a pioneer transcription factor, a specialized subclass of transcription factors that can bind nucleosomal DNA prior to generation of local accessibility. However, evidence at the genome scale for PAX3/7::FOXO1 direct nucleosome targeting, prior to the accessibility step in pioneering, has remained elusive and challenging to capture methodologically for RMS fusion oncoproteins. In this work, we compare the cellular functions of these PAX::FOXO1 fusions, including new approaches for identifying nucleosome targeting at the genome scale. We find that in zebrafish RMS initiation models, the fusions initially activate similar neural transcriptional programs but to different extents, and we further evaluate their mechanisms in RMS cells at the genome scale with modified MNase XChIP to detect nucleosome and subnucleosome fusion/chromatin binding. In establishing our cross-species comparative oncology approach, we report, to our knowledge, the first high resolution nucleosome positioning data in rhabdomyosarcoma. We find that both PAX::FOXO1 fusions bind nucleosomal DNA, but with varied motif preferences and histone mark co-localization patterns. Altogether, we establish the nucleosome targeting functions of PAX7::FOXO1 and PAX3::FOXO1 pioneering and uncover key mechanistic distinctions for chromatin engagement of the two most common RMS fusion oncoproteins.

**HIGHLIGHTS:**

- Partially overlapping gene signatures are activated by PAX3/7::FOXO1 in vivo
- Modified MNase ChIP reveals PAX3/7::FOXO1 bind nucleosomal and subnucleosomal DNA
- PAX7::FOXO1 binds degenerate paired/homeobox motifs within nucleosome targets
- Each fusion engages distinct nucleosomal gene targets

## INTRODUCTION

Rhabdomyosarcoma (RMS) is an aggressive pediatric cancer and the most common soft tissue sarcoma in children^1^. Although historically categorized based on histology, more recent and clinically predictive approaches have focused on the presence or absence of a fusion oncoprotein^2–4^. Despite decades of research, outcomes for many patients with higher risk factors like *PAX*-fusion status remain poor^5^. Additionally, the standard of care for treating young RMS patients still includes broadly cytotoxic chemotherapies, surgery, and radiation^6^. These treatments are often not curative and can cause lifelong adverse side effects^7^. However, there is increasing evidence for epigenetic drivers of pediatric tumors, including RMS^8–12^. Therefore, there is a pressing need for a better understanding of the central epigenetic mechanisms of this cancer and the development of targeted therapeutics are critically needed, as epigenetic processes are reversible.

RMS fusions are often caused by a reciprocal chromosomal translocation event and drive a specific subtype of this disease, called fusion-positive rhabdomyosarcoma (FP-RMS). The most common fusions are between either *PAX3* (chr. 2q) or *PAX7* (chr. 1p) fused to *FOXO1* (chr. 13q)^13–15^. Tumorigenic PAX::FOXO1 fusions contain both a paired and a homeodomain DNA binding domain (DBD) from PAX3/PAX7 as well as a partial forkhead DBD and full transactivation domain from FOXO1. Wild-type PAX3 and PAX7 are key transcription factors involved in myogenic and neural crest development with distinct, partially redundant roles^16–18^. As such, with 86% sequence similarity, Pax7 can substitute for Pax3 in neural crest and somite functions, but not within muscle satellite cells^18^. Wild-type FOXO1 is involved in cellular metabolism processes, including gluconeogenesis and adipogenesis, and resistance to cellular stress^19^.

Both PAX3 and PAX7 have certain features consistent with pioneer function, likely arising from their PAX3/7 DNA binding domains (DBDs; preserved in the major RMS fusion oncoproteins), where they can interact with condensed, mitotic, or closed chromatin and induce accessibility changes^20–26^. The PAX3::FOXO1 fusion, which preserves the intact PAX3 DBD, can also engage closed chromatin and induce local accessibility^8,27,28^. However, the chromatin engagement mechanisms of PAX7::FOXO1 have not been extensively studied. Despite the shared histological features between PAX3::FOXO1 and PAX7::FOXO1-positive tumors, the presence of PAX3::FOXO1 is associated with more aggressive disease and worse patient outcomes than PAX7::FOXO1^29,30^. These shared features and disparities in patient outcomes provide motivation to uncover the epigenetic mechanisms driving differences in aggressiveness of PAX3::FOXO1 and PAX7::FOXO1 as fusion drivers of RMS. While recent advances have revealed key functional differences between these fusion proteins within in vitro exogenous over-expression models and IPSC-derived myogenic precursor cells^31,32^, the precise mechanisms for chromatin invasion as an epigenetic antecedent to inducing enhancer-promoter interactions remain elusive. Here, we sought to epigenetically and functionally compare PAX3::FOXO1 and PAX7::FOXO1 using a cross-species approach in vertebrate zebrafish embryos and patient-derived cell lines. Herein, we compare the initial transcriptional profiles driven by each fusion protein in vivo. Additionally, we developed a modified MNase XChIP approach to investigate mechanistic differences in nucleosomal motif preferences and binding in cell-based studies.

## RESULTS

### PAX::FOXO1 fusions drive similar initial transcriptional signatures in zebrafish

Pioneer factors can target inactive or nucleosomal regions of the genome to anticipate or reprogram cellular identities^33–40^. We were motivated to understand any functional differences in the initial programs that PAX3::FOXO1 and PAX7::FOXO1 fusions were activating during early steps in epigenetic reprogramming. To determine early-immediate functional activities of the fusions, we developed a zebrafish *PAX7::FOXO1* mRNA injection model for comparative analysis to PAX3::FOXO1 (**Figure 1A**), conceptually similar to our recently reported approach^28,41^. Our in vivo work with PAX3::FOXO1 zebrafish models recently revealed that the human fusion initially activated neural transcriptional programs present in the human disease^28^. We injected equimolar amounts of *PAX3::FOXO1* or *PAX7::FOXO1* mRNA and observed that *PAX7::FOXO1* injection resulted in a slight, albeit statistically significant (*p* = 0.012) improvement in overall embryo survival (**Supplemental Figure S1A-C**). We observed that the fusions drive partially overlapping transcriptional programs, with key functional distinctions (**Figure 1B**). For example, when compared to control-injected embryos, PAX7::FOXO1 expression resulted in 6,534 upregulated genes versus 5,386 for PAX3::FOXO1 (**Supplemental Figure S1D**). Moreover, PAX7::FOXO1 activated gene expression to a greater magnitude increase than PAX3::FOXO1 (**Supplemental Figure S1E**). Altogether, differential expression analysis revealed that PAX7::FOXO1 is a stronger initial transcriptional activator, in agreement with its function when ectopically expressed in human fibroblasts^31^.

**Figure 1.**
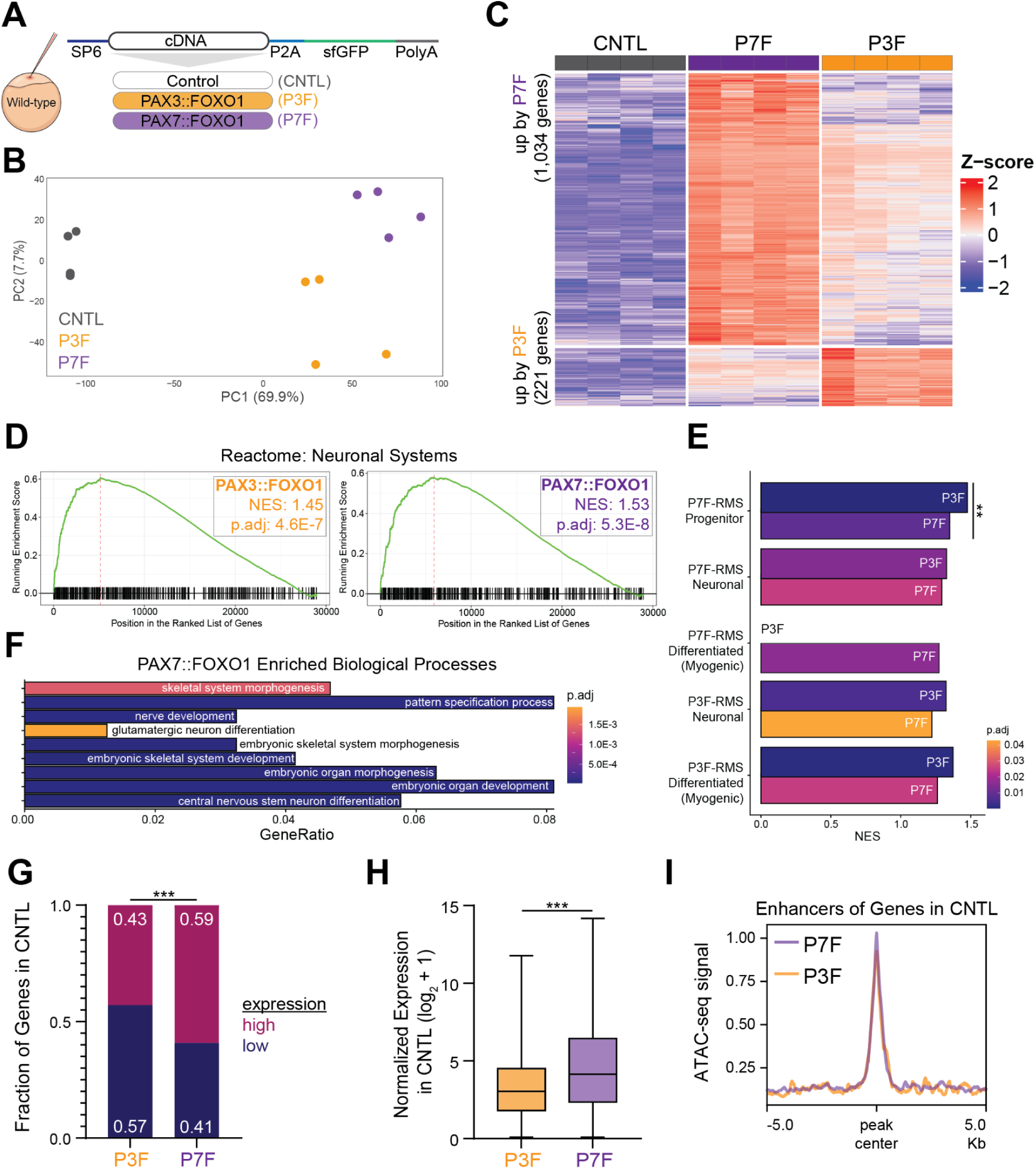
PAX3::FOXO1 and PAX7::FOXO1 initially induce similar transcriptional programs in zebrafish embryos but have key distinctions in gene targeting. (A) Wild-type zebrafish embryos were injected at the one-cell stage 100 ng/μL of PAX3::FOXO1 (P3F) mRNA or equimolar PAX7::FOXO1 (P7F) or Control (CNTL) mRNA. (B) PCA of RNA-seq in CNTL, P3F, and P7F-injected embryos. (C) Z-score normalized heatmap of differential activated genes between PAX::FOXO1 fusions versus CNTL-injected embryos. (D) Gene set enrichment analysis (GSEA) to neuronal systems reactome pathway for P3F or P7F versus CNTL. (E) Gene over-representation analysis (ORA) to fusion-positive rhabdomyosarcoma markers on the y-axis, identified from scRNA-seq analysis in Danielli et al., 2024, for P3F or P7F versus CNTL embryos, in bars. Asterisks represent if statistically significant from ORA between P3F and P7F embryos. Statistics show if markers are enriched in differentially expressed genes in Figure 1C. (*) indicates 0.01 > *p* > 0.001. (F) ORA to biological processes enriched in genes upregulated by P7F in Figure 1C. (G) Fraction of genes in (C) that are highly or lowly expressed in CNTL embryos. Normalized reads of 10 was the cutoff for high/low expression. A Fisher’s exact test was used to determine statistical significance. (H) Violin plot of log2+1 normalized expression of genes from Figure 1C in CNTL embryos. A Mann-Whitney test was used to determine statistical significance. (I) Profile plot of average ATAC-seq signal for the enhancers regulating genes from Figure 1C in CNTL embryos. Enhancer-gene interactions were determined according to the activity-by-contact model from Kucinski et al., 2025. (**) indicates 0.01 > *p* > 0.001, and (***) indicates *p* < 0.001.

Previously, we showed that PAX3::FOXO1 expression in embryonic zebrafish results in transcriptional activation within hours with gene signatures that are consistent with RMS patient gene signatures^28^. Given this, we focused on genes differentially activated between the PAX3/7::FOXO1 fusion oncoproteins on rapid timescales. In terms of immediate-early transcriptional targeting and direct comparison between fusions, we observed that PAX7::FOXO1 more strongly activated a greater number of genes than PAX3::FOXO1 (1,034 versus 221) (**Figure 1C**). However, a subset of those genes was also rapidly upregulated by PAX3::FOXO1, although to a lesser extent. Gene set enrichment analysis (GSEA) revealed that both fusions rapidly upregulated similar transcriptional programs, including neuronal system, synaptic, and GPCR signaling (**Figure 1D; Supplemental Figure S1F**). Examining overlap, 60.9% of activated biological processes by PAX7::FOXO1 were also activated in PAX3::FOXO1-injected embryos (**Supplemental Figure S1G-H**). Moreover, each fusion activated markers for similar FP-RMS cell states that were previously defined from single-cell data (**Figure 1E**)^42^. Lastly, comparison between fusions showed that PAX7::FOXO1 activated neural and broad developmental processes (**Figure 1F**), whereas PAX3::FOXO1 better activated progenitor population markers from PAX7::FOXO1 tumors (**Figure 1E**).

We were intrigued that, despite PAX7::FOXO1 rapidly inducing stronger gene activation, its expression also resulted in less severe phenotypic consequences than PAX3::FOXO1. This observation led us to hypothesize that contrasting capabilities of the fusions to bind to and activate closed chromatin could influence epigenetically divergent outcomes. Towards this end, we observed that genes more strongly activated in PAX7::FOXO1-injected embryos were more often already activated in our control embryos. We reasoned that there might be epigenetic priming of key gene targets prior to PAX7::FOXO1 activity. In alignment with this conceptual framework, 57.1% of PAX3::FOXO1-enriched genes were lowly expressed (less than 10 normalized reads) compared to 40.9% of PAX7::FOXO1-enriched genes (**Figure 1G**). PAX7::FOXO1-enriched genes were also more highly expressed in the control embryos, with their enhancers having more accessibility in controls (**Figure 1H-I**). Overall, our studies with fusion inductions in zebrafish reveal that PAX::FOXO1 fusions rapidly activate similar programs, despite PAX7::FOXO1 being a stronger activator, and suggest that differences in their mechanisms of invasion of closed chromatin may critically factor into phenotypic outcomes.

### PAX::FOXO1 fusions cellularly localize to both accessible and inaccessible chromatin

Since pioneer factors can interact with closed chromatin, we utilized biochemical approaches to determine if PAX7::FOXO1 can reside in several distinct nuclear fractions, or if it is primarily localized within the soluble nucleoplasm. Using established RMS patient cell lines endogenously expressing PAX7::FOXO1 (CW9019), PAX3::FOXO1 (RH4), and a fusion negative patient cell line (SMS-CTR), we sequentially isolated cellular proteins via sonication-based and salt-based cellular fractionation methods^37,43,44^. With sonication-based fractionation, isolated nuclei are further resolved via mechanical shearing; soluble-euchromatin is sensitive to sonication, and insoluble-heterochromatin is resistant to sonication^37,45^. For the salt-based fractionation, increasing salt concentrations causes a sequential release of nuclear proteins, demarcating euchromatin in lower salt concentrations and heterochromatin in higher salt concentrations^46^.

We confirmed the effectiveness of these protocols to progressively extract proteins associated with varying levels of chromatin accessibility by concurrently identifying the localization of BAF155, TBP, and HP1. BAF155, encoded by *SMARCC1*, is a subunit of the mammalian SWI/SNF ATP-dependent chromatin remodeling complex, and is found enriched in the nuclear and soluble chromatin fractions by the sonication-based protocol and the 250mM and 600mM NaCl fractions by the salt-based protocol (**Figure 2A,B**). HP1 reads and nucleates the heterochromatin histone modification H3K9me3 and has critical roles in the establishment and maintenance of silenced heterochromatin. In the sonication fractionation, HP1 (α/β) has the strongest bands within the insoluble chromatin pellet fraction, while there is also some signal in the soluble nuclear and chromatin fractions, and it is localized mainly to the 250 mM NaCl and 600mM NaCl fractions in the salt-based fractionation (**Figure 2A,B**). These patterns are consistent with a repressive chromatin adaptor protein directly associating with and reinforcing local inaccessible chromatin states. TBP has ubiquitous expression and functions during gene activation and mitotic bookmarking, and was used to confirm and compare loading within each nuclear fraction (**Figure 2A,B**)^47^. We blotted for epitopes preserved in PAX::FOXO1 fusion proteins with an antibody for the C-terminus of FOXO1^43,48,49^ and observed strong signal for both fusion oncoproteins, as well as evidence for cytoplasmic FOXO1 monomer in the fusion-negative context. Consistent with previous findings, expression of both PAX::FOXO1 fusions resulted in substantial downregulation of wild-type FOXO1 protein compared to fusion-negative RMS (SMS-CTR), and wild-type FOXO1^43,50^. In contrast, PAX3::FOXO1 and PAX7::FOXO1 were increasingly seen in both euchromatic and heterochromatic fractions, highlighting a capability to localize within different biochemical and regulatory chromatin regions, a capacity which we hypothesize enables target gene activation from reduced expression states (**Figure 2A,B**).

**Figure 2.**
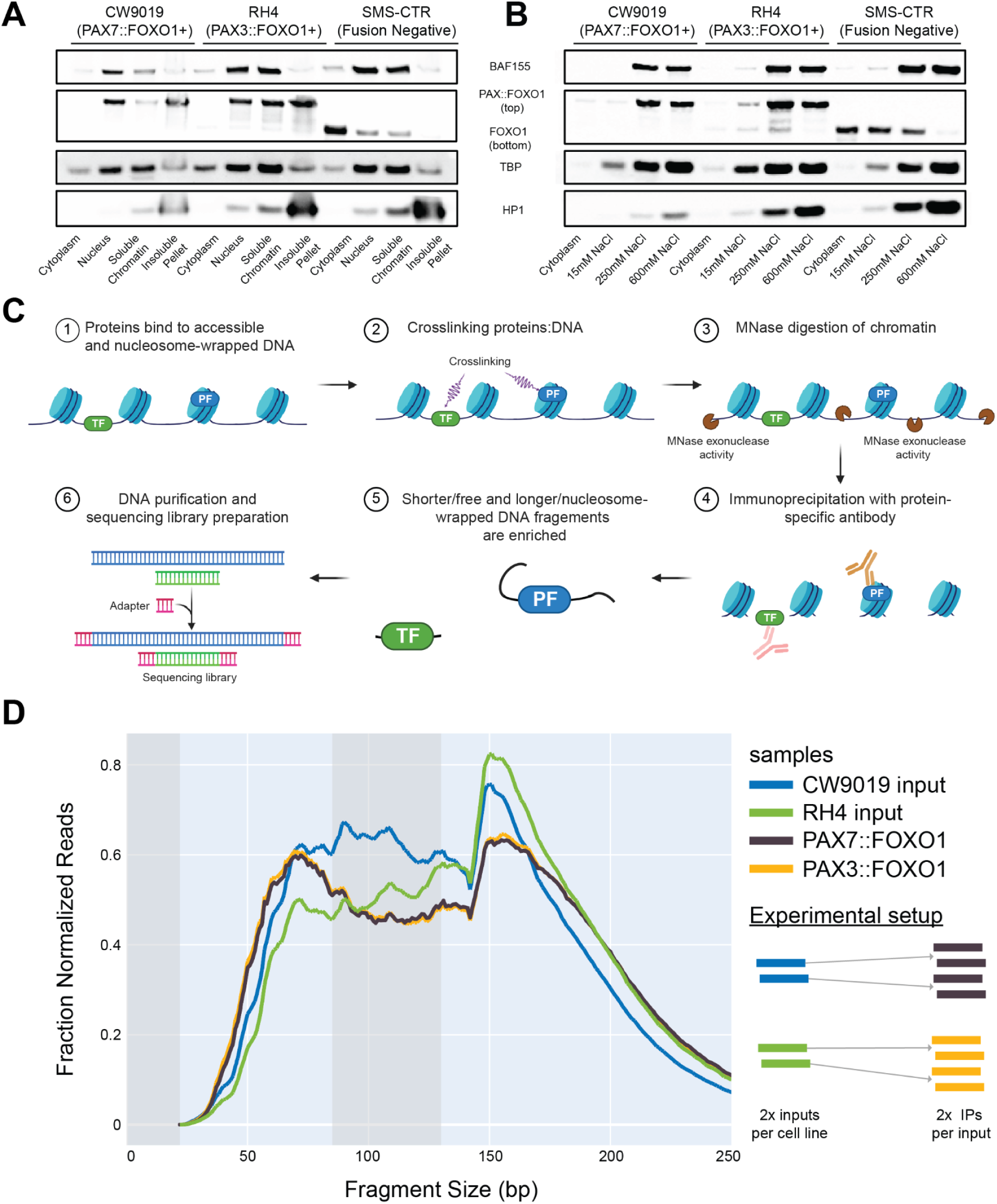
PAX3::FOXO1 and PAX7::FOXO1 localize within nucleosome-enriched regions and closed chromatin regions in addition to open chromatin with cellular fractionation and modified MNase XChIP methodologies. (A-B) Western blot for sonication-based and salt-based fractions, respectively, of CW9019 (PAX7::FOXO1 expressing), RH4 (PAX3::FOXO1 expressing), and SMS-CTR (fusion negative, FOXO1 expressing) rhabdomyosarcoma patient cell lines. (C) Workflow for modified MNase XChIP. TF denotes transcription factor and PF denotes pioneer factor. (D) Distribution of modified MNase XChIP fragment length. Averages of all replicates are shown, including 2 input and 4 anti-PAX::FOXO1 immunoprecipitation replicates per cell line. Intermediate fragment lengths (86-130 bp) excluded from downstream analysis are shown in gray.

### Modified MNase XChIP enables simultaneous genome-wide identification of nucleosomal and nucleosome-free DNA binding by pioneer factors

While there is evidence that PAX3::FOXO1 can induce local DNA accessibility, or nucleosome depletion, at its target sites, we were motivated to investigate more direct evidence for nucleosome targeting which has remained elusive^28,43,51^. Moreover, evidence for pioneering activity has not been assessed for PAX7::FOXO1, while there is evidence for its functional capacity in transcription activation^31,32^. Therefore, we sought to develop a strategy to investigate the fusions’ engagement with both nucleosomal (pioneering sites) and subnucleosomal (accessible sites) chromatin at the genome scale. To simultaneously capture sequencing data containing both types of interactions while preserving the ability to distinguish between them, we modified a previously published MNase-based approach, MNase XChIP (**Figure 2C**)^52^. MNase XChIP was originally designed to improve the resolution of canonical TF-DNA interactions similar to ChIP-exo footprinting studies^53^. Our modifications enable the capture of TF-nucleosome interactions, giving us the opportunity to measure chromatin engagement loci as well as epigenetic context within the same experiment.

In our modified MNase XChIP, chromatin-interacting proteins, such as TFs (or pioneer TFs), chromatin regulatory complexes, and nucleosomes are crosslinked to DNA and digested with MNase (see **Methods**). The exo- and endo-nuclease activities of MNase digest exposed DNA, while protein-bound DNA is protected from digestion. The core length of DNA protected by being part of a nucleosome is ∼147 bp, and TFs provide a smaller footprint of protected DNA^52,54^. Immunoprecipitation (IP) identifies protein-DNA interactions, with input controls identifying nucleosome occupancy patterns from the MNase (excluding IP). Thus, input controls can also serve as a non-size-selected MNase-seq capturing nucleosome positioning and occupancy information^55,56^. By removing size-selection steps during library preparation, we can preserve nucleosomal and TF-footprint-sized DNA fragments.

We applied this modified methodology independently to either PAX7::FOXO1-expressing or PAX3::FOXO1-expressing RMS patient cell lines (CW9019 and RH4, respectively), with two MNase digests per cell line and two PAX::FOXO1 (anti-FOXO1) IPs per digest, extending our ChIP-seq approach^28,41,43^ for other fusions in RMS. The fragment length distributions for PAX3::FOXO1- and PAX7::FOXO1-bound chromatin are remarkably consistent, with a clear bimodal distribution of nucleosome-sized fragments and subnucleosomal PAX::FOXO1-bound fragments (**Figure 2D**). Nucleosomal MNase-seq (MNase XChIP input) fragments have the highest read densities at 151 bp, with a steep increase in relative fraction of reads beginning at 142 bp. Notably, the PAX3::FOXO1 and PAX7::FOXO1 MNase XChIP IP samples also have the highest read densities near the nucleosomal fragment length, although they have a wider distribution at longer lengths than the MNase-seq controls (input samples). This pattern is consistent with nucleosome engagement by the fusion proteins protecting additional adjacent DNA to the core of nucleosome-wrapped DNA. In subnucleosomal DNA fragments from our MNase-seq, there are less distinct maxima in read densities, which may reflect cell type-specific patterns. Both PAX3::FOXO1 and PAX7::FOXO1 MNase XChIP IPs have a single subnucleosomal distribution centered at approximately 70 bp, consistent with fragment lengths seen in MNase XChIP studies of CTCF^52^. Overall, the bimodal distribution of nucleosomal and subnucleosomal fragments in PAX3::FOXO1 and PAX7::FOXO1 MNase XChIP IPs reveal that these fusion proteins can bind both nucleosome-wrapped and exposed DNA.

We sorted each sample’s reads into either nucleosomal or subnucleosomal bins *in silico* and analyzed these categories separately. To preserve the majority of sequencing reads, we tested different cutoffs for binning based on fragment length and compared intermediate fragments nucleosomal and subnucleosomal categories (<90 and ≥124 bp and <85 and ≥130 bp). Reads between nucleosomal and subnucleosomal distributions in our data may be representative of partially unwrapped digested nucleosomes^57^, hexasomes that lack one H2A/H2B dimer from the canonical nucleosome octamer and therefore only occupy ∼110-115bp of DNA^58^, or hemisomes/half-nucleosomes that consist of a single copy of each histone^59,60^. We focused our analyses on more clearly distinguished and identifiable nucleosomal (defined as ≥130 bp) and subnucleosomal (<85 bp) fragment length populations (**Figure 3A**).

**Figure 3.**
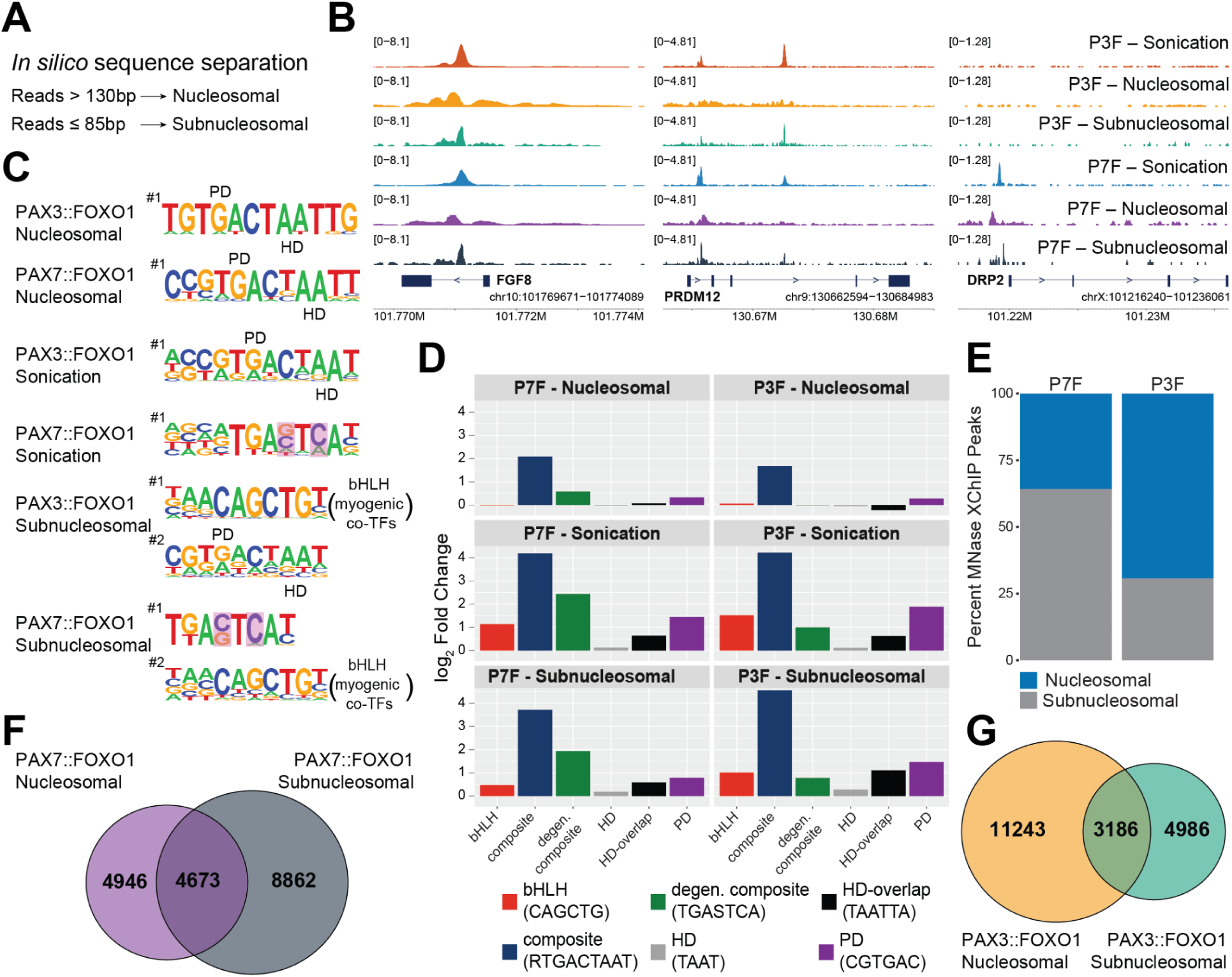
Nucleosome-fusion protein binding for both PAX3::FOXO1 and PAX7::FOXO1 in RMS cells is associated with the higher complexity composite paired-homeobox motif. (A) *In silico* fragment length sorting strategy for nucleosomal and subnucleosomal bins. (B) Nucleosomal and subnucleosomal modified MNase XChIP and sonication ChIP signal tracks at example loci *FGF8* (left), *PRDM12* (middle), and *DRP2* (right), for PAX3::FOXO1 (P3F) and PAX7::FOXO1 (P7F). (C) Top enriched *de novo* consensus binding motifs for each peak set. Portions of each motif are marked corresponding to the paired domain (PD) and homeobox domain (HD) motifs, the motifs for the two DNA-binding domains present in the PAX3 and PAX7 transcription factors. Variation at key bases is highlighted in pink for P7F sonication and subnucleosomal motifs. The top two motifs are shown for subnucleosomal peak sets to show enrichment for the basic helix-loop-helix (bHLH) motif bound by myogenic regulatory factors known to interact with PAX::FOXO1 proteins. (D) Calculated log_2_ fold-change enrichments for PD, HD, overlapping homeobox (HD-overlap), composite (PD-HD), degenerate composite, and bHLH motifs within each peak set compared to background. (E) Comparison of percentages of unique nucleosomal to subnucleosomal peaks for P7F and P3F. (F-G) Venn diagrams showing overlap of nucleosomal and subnucleosomal peaks for P7F and P3F, respectively.

### Nucleosomal MNase XChIPs exhibit distinct binding patterns and motif preferences compared to subnucleosomal or sonication-based ChIPs

When visually comparing sequencing results, *subnucleosomal* PAX3::FOXO1 and PAX7::FOXO1 MNase XChIP IPs largely resemble their respective sonication-based ChIPs. In contrast, in silico separated *nucleosomal* MNase XChIP IP profiles for both fusion oncoproteins appear distinct from both *subnucleosomal* and *sonication* profiles, and reveal binding that can mirror nucleosome phasing and periodicity (see *FGF8* locus; **Figure 3B**, left)^61,62^. However, there are also key examples of distinct binding patterns between PAX3::FOXO1 and PAX7::FOXO1. When comparing PAX3::FOXO1 and PAX7::FOXO1 binding patterns at the *PRDM12* gene locus, PAX7::FOXO1 has increased binding towards the TSS, and PAX3::FOXO1 binds within the gene body in both subnucleosomal MNase XChIP profiles and sonication ChIP-seq profiles (**Figure 3B**, middle). Another example of divergent binding activity between PAX3::FOXO1 and PAX7::FOXO1 is at the *DRP2* gene locus, where high PAX7::FOXO1 binding occurs near its TSS and PAX3::FOXO1 has only very weak, background-level signal throughout, consistent across both MNase XChIP and ChIP-seq approaches (**Figure 3B**, right). Together, our results highlight key differences in chromatin binding profiles between PAX3::FOXO1 and PAX7::FOXO1 fusion oncoproteins. These results are significant for the field, suggesting that there are likely distinct core regulatory circuitries for each fusion oncoprotein.

Enriched DNA binding motifs were identified *de novo* using HOMER on *in silico* separated consensus peak sets for nucleosomal, subnucleosomal, and sonication ChIPs for each fusion (**Figure 3C**)^63^. For both PAX3::FOXO1 and PAX7::FOXO1 binding, the top enriched DNA-binding motif within nucleosomes was for the composite PAX3/PAX7 motif^22,43^, consisting of a combination of the motifs bound by paired domain (PD) and homeodomain (HD) DNA binding domains preserved in the fusion oncoproteins. PAX7::FOXO1 was also found to bind to this composite motif in iPSC-derived myogenic precursor cells^32^. Our most enriched consensus motifs for PAX3::FOXO1 binding peak sets from sonication ChIP-seq and subnucleosomal MNase XChIP very closely match previous findings, with enrichments for a composite PD-HD motif, while there was some increased variation from the precise bases seen in the nucleosomal *in silico* separated sample^43,64^. However, the PAX7::FOXO1 sonication (ChIP-seq) and subnucleosomal peak sets from MNase XChIP have even more variation from the canonical bases of the composite PD-HD motif, particularly at the final base in the PD portion (C to C/G) and second base of the HD (A to C) (**Figure 3C**, highlighted bases). Due to its key differences to the canonical composite PD-HD motif, we refer to this motif as a degenerate composite. It also has a high degree of sequence similarity to those bound by bZIP TFs^43^. Additionally, the top motif for PAX3::FOXO1 subnucleosomal (MNase XChIP) peaks is an E-box motif, specifically from a bHLH TF family that includes known myogenic regulatory factors (MRFs) MYOD, MYOG, MYF5^51,65^. The same motif is the second most enriched in the PAX7::FOXO1 subnucleosomal (MNase XChIP) peak set.

We used these same peak sets from our MNase XChIP and sonication-based ChIP-seq to compare the relative enrichments of key known DNA motifs based on those previously identified for PAX3, PAX7, and their respective PAX::FOXO1 fusions^22,31,64^. As is consistent with the *de novo* identified motifs, this degenerate composite PD-HD motif is particularly enriched in PAX7::FOXO1’s sonication ChIP-seq and subnucleosomal MNase XChIP peaks (log2 fold-changes of 2.4 and 1.9, respectively), and it is only somewhat enriched at PAX3::FOXO1’s sonication ChIP-seq and subnucleosomal MNase XChIP (0.99 and 0.78) sites and not at all in nucleosomal peaks (-0.01) (**Figure 3D**). Another striking pattern is the co-localization with bHLH motifs bound by MRFs. This enrichment is higher for subnucleosomal PAX3::FOXO1 in comparison to PAX7::FOXO1 MNase XChIP targets, but is nearly absent in both nucleosomal MNase XChIP peak sets and may therefore indicate relatively decreased cooperativity with MRFs in these contexts. Additionally, while sonication ChIP-seq and nucleosomal MNase XChIP peaks for both fusions are enriched for the canonical PD and an *overlapping palindromic* HD motifs, this is not seen in the nucleosomal categories, which are distinctly enriched for the composite PD-HD motif. Overall, sonication ChIP-seq and subnucleosomal MNase XChIP peaks again appear largely similar to each other, with nucleosomal binding by both PAX::FOXO1 fusions being strongly associated with a more precise version of the PD-HD composite motif.

For each fusion, we determined the overlaps between called nucleosomal, subnucleosomal, and sonication consensus peaks (from MNase XChIP and ChIP-seq, respectively) and observed that the largest overlaps for each fusion are between their respective subnucleosomal and sonication peak sets (**Supplemental Figure S3**). PAX7::FOXO1 has substantially more overlaps across its peak sets, whereas PAX3::FOXO1 nucleosomal MNase XChIP peaks are the largest and most distinct set. Furthermore, we observed that a higher fraction of PAX3::FOXO1 MNase XChIP peaks were in nucleosomal DNA, suggesting differential capabilities between the respective fusion proteins to bind nucleosomes (**Figure 3E**). Direct pairwise comparisons between nucleosomal and subnucleosomal MNase XChIP peaks for each fusion oncoprotein revealed that PAX7::FOXO1 has a substantially higher relative proportion of peaks shared between nucleosomal and subnucleosomal (**Figure 3F,G**). This could be indicative of PAX7::FOXO1 having a more similar mode of nucleosomal and subnucleosomal binding and/or a higher rate of nucleosome turnover at its binding sites. More rapid nucleosome turnover would result in an increased likelihood of binding motifs occurring on exposed or partially accessible DNA. In summary, PAX3::FOXO1 has a greater propensity to bind nucleosomal DNA than PAX7::FOXO1 and appears to bind to those sites through recognition of a precise PD-HD motif instead of a degenerate composite motif.

### PAX7::FOXO1 and PAX3::FOXO1 have divergent modes of interacting with well-positioned nucleosomes

We next wondered if our MNase XChIP approach for capturing the chromatin context for fusion oncoprotein binding events could reveal any additional differences in how the PAX::FOXO1 proteins interact with nucleosomes. We plotted all MNase XChIP reads at nucleosomal and subnucleosomal peaks as well as sonication ChIP-seq peaks for both PAX7::FOXO1 and PAX3::FOXO1 (**Figure 4A,D**). Peaks were further filtered to only include those with putative composite PD-HD motifs for binding, while allowing for one mismatch from the canonical central portion of the motif. Reads were plotted using plot2DO (ref.^66^) at regions 500bp upstream and downstream from the potential binding motifs within peaks (**Figure 4A,D**, x-axis). Reads are further characterized by fragment length (**Figure 4A,D**, y-axis) and relative coverage of reads for each set of regions were represented colorimetrically in heatmaps. As expected, sonication-based ChIP-seq for both PAX7::FOXO1 and PAX3::FOXO1 displayed a range of fragment lengths centered on the motif, consistent with random shearing of chromatin via sonication yielding a range of DNA fragment sizes (**Figure 4A,D**, bottom). Subnucleosomal PAX7::FOXO1 and PAX3::FOXO1 (**Figure 4A,D**, middle) also showed binding centered around the motif, with fragment length distributions consistent with the footprints protected from MNase digestion by PAX3::FOXO1 or PAX7::FOXO1 binding of nucleosome-free DNA (**Figure 4A,D**, middle).

**Figure 4.**
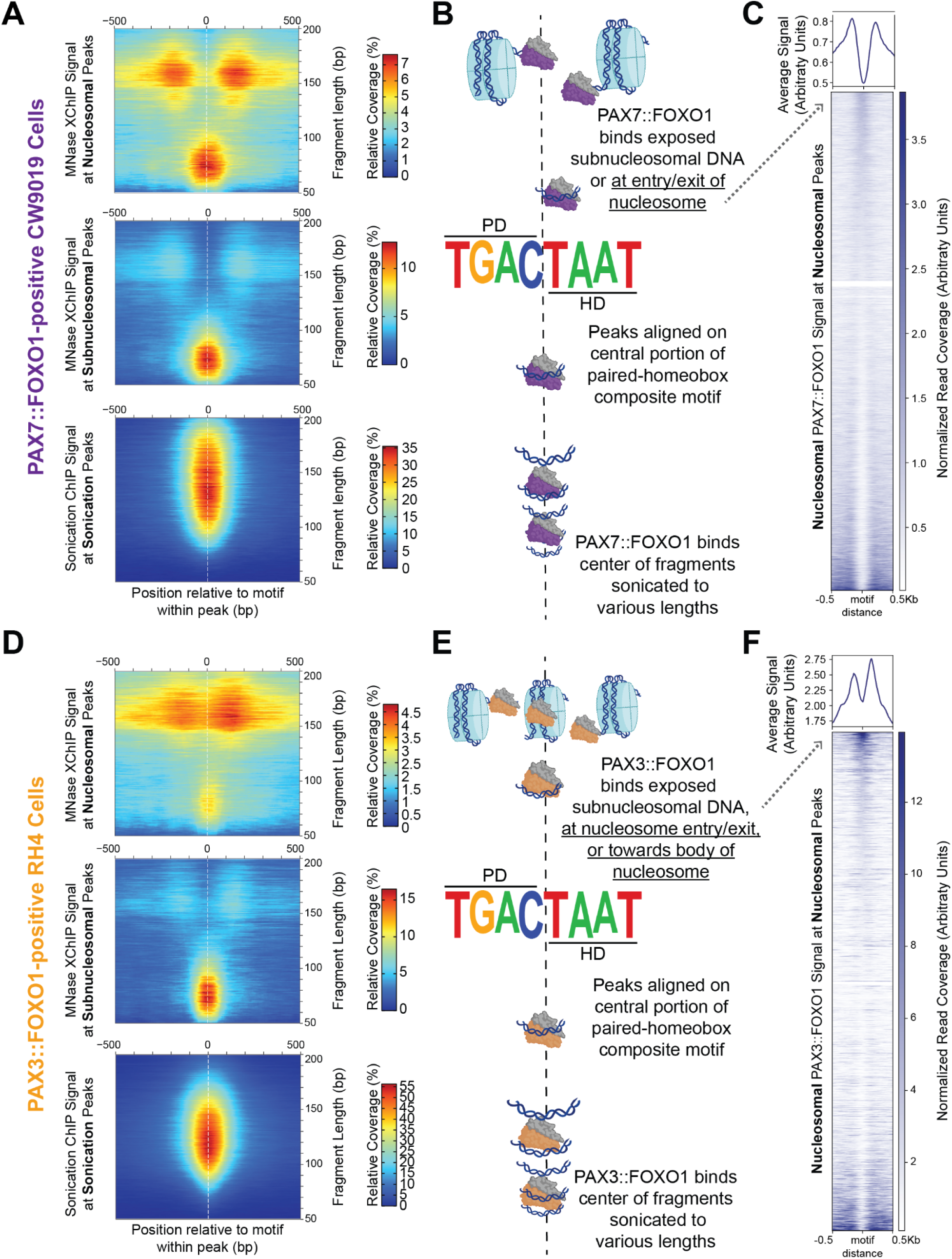
PAX3::FOXO1 and PAX7::FOXO1 have distinct chromatin binding modes at well-positioned nucleosomes in RMS cells. (A) PAX7::FOXO1 MNase XChIP signal at nucleosomal peaks (top), at subnucleosomal peaks (middle), and sonication ChIP signal at sonication peaks (bottom). Peak regions are centered on PD-HD target motifs (x-axis) with sequencing reads at these regions separated based on fragment length plotted (y-axis). The relative read coverage is shown by the heatmap color intensity (colorimetric scale insets). (B) Representations of PAX7::FOXO1 chromatin binding mechanisms corresponding to adjacent heatmaps. The DNA-binding motif used for centering data from heatmaps, (allowing for 1 mismatch) is shown. (C) Profile plot (top) and heatmap (bottom) isolating the nucleosomal signal across the same regions shown (A, top). (D) PAX3::FOXO1 MNase XChIP signal at nucleosomal peaks (top), at subnucleosomal peaks (middle), and sonication ChIP signal at sonication peaks (bottom). Peak regions are centered on PD-HD target motifs (x-axis) with sequencing reads at these regions separated based on fragment length plotted (y-axis). The relative read coverage is shown by the heatmap color intensity (colorimetric scale insets). (E) Representations of PAX3::FOXO1 chromatin binding mechanisms corresponding to adjacent heatmaps. The DNA-binding motif used for centering data from heatmaps, (allowing for 1 mismatch) is shown. (F) Profile plot (top) and heatmap (bottom) isolating the nucleosomal signal across the same regions shown (D, top).

However, differences between the fusions become apparent when comparing read length and positioning at nucleosomal PAX7::FOXO1 and PAX3::FOXO1 peak regions (**Figure 4A,D**, top). We found that nucleosomal binding events at fragment lengths above ∼150 bp for PAX7::FOXO1 are centered around 150 bp from binding motifs. We interpret these data to be the result of PAX7::FOXO1 binding its PD-HD motif at or near the entry/exit site of nucleosome-wrapped DNA, with the MNase cleavage reaction occurring +/- 150 bp depending on the direction of nucleosome positioning relative to the motif. While PAX3::FOXO1 also appears to be able to target the entry/exit site of the nucleosome, there is a substantially less clear gap or distinction between the two populations (**Figure 4D**, top). We interpret this to be due to an additional capability for PAX3::FOXO1 to bind motifs that are positioned further within the body of the nucleosome. It should also be noted that nucleosomal PAX7::FOXO1 peaks show prominent subnucleosomal binding (**Figure 3F**, **Figure 4A**, top). The higher overlap of PAX7::FOXO1’s nucleosome and subnucleosomal peak regions (MNase XChIP, **Figure 3F**) compared to PAX3::FOXO1 may also result from higher nucleosome turnover at its +1 nucleosome targets, or partial destabilization as PAX7::FOXO1 targets entry/exit sites (**Figure 4A,D**)^67^. For each set of plots, our proposed models to explain these binding interaction data are shown in **Figure 4B,E**. To further clarify differences for nucleosomal binding alone, we plotted nucleosomal PAX7::FOXO1 and PAX3::FOXO1 binding signal (**Figures 4C,F** profile plots) and sorted by intensity within the central 150 bp (**Figures 4C,F** heatmaps, top) or outside the central 150bp (**Figures 4C,F** heatmaps, bottom). These data confirm the overall preference for binding at entry/exit sites for both fusions, while PAX3::FOXO1 also has notable binding in the central nucleosome-sized region (**Figure 4C,F**).

### Nucleosomes downstream of actively transcribed genes are enriched for both PAX3::FOXO1 and PAX7::FOXO1 fusions

Given the differences in PAX3::FOXO1 and PAX7::FOXO1 nucleosomal targeting, we also questioned if there were functional differences in gene expression related to these binding modes at +1 nucleosomes bound by each fusion. When examining read density at protein-coding gene regions (**Supplemental Figure S3**, center), both sonication ChIP-seq and *in silico* separated subnucleosomal MNase XChIP data showed similar trends with increasing read density upstream of TSSs, followed by sharp decreases in read density within gene bodies. MNase XChIP subnucleosomal input data displayed increased noise, which may be driven from rapid on and off rates of chromatin regulatory machinery within the genome. In contrast, *in silico* separated nucleosomal MNase XChIP inputs had a general depletion within gene bodies and sharp decreases at TSSs and TESs, corresponding to 5’ and 3’ nucleosome-free regions (NFRs, ref.^61^). Interestingly, both PAX7::FOXO1 and PAX3::FOXO1 nucleosomal reads from our MNase XChIP data have a second peak immediately downstream of the TSS within the gene body, which we interpret to be bound +1 nucleosomes. To visualize these interactions, we plotted both sonication ChIP-seq data and MNase XChIP data using plot2DO^66^ within 1 kb up or downstream of the same gene loci (**Supplemental Figure S3**). Again, sonication ChIP-seq showed few differences, with uniform distribution of fragment lengths centered on TSSs for PAX3::FOXO1, PAX7::FOXO1 and input samples (**Supplemental Figure S3**, ChIP-seq occupancy plots). Our MNase X-ChIP input (MNase-seq) data showed strong +1 nucleosome and 5’ NFR signals, and there were few subnucleosomal reads for MNase XChIP inputs, which is reflected in the **Supplemental Figure S3** occupancy plots. To our knowledge, this is the initial report of nucleosome positioning data in RMS.

Both PAX7:FOXO1 and PAX3::FOXO1 MNase XChIPs show similar features in their occupancy plots, with strong nucleosomal binding at the +1 nucleosome. The subnucleosomal MNase XChIP enrichments centered in proximity with TSSs mirror from the observations from our sonication ChIP-seq data. In both cases, PAX7::FOXO1 has notably stronger enrichments for each of these features relative to PAX3::FOXO1, potentially indicating more prevalent or stronger interactions at gene promoters, which has also been observed in another recent report^32^. Overall, the fusions’ engagement with +1 nucleosomes across the genome, in addition to PAX3::FOXO1 regulation of nascent chromatin, suggests PAX::FOXO1 fusions may play key roles in temporal regulation of nucleosome phasing and epigenome organization^68^.

### PAX::FOXO1 fusions have contrasting associations to nucleosomes within local regions of active and repressive histone marks

Having established nucleosomal binding patterns for PAX3::FOXO1 and PAX7::FOXO1 fusions, as well as obtaining information on well-positioned nucleosomes in RMS, we sought to place fusion/nucleosome engagement events within the context of surrounding histone post-translational modifications (PTMs). We meta-analyzed our MNase XChIP data for each fusion with sonication-based ChIP-seq in each cell line for histone PTMs commonly associated with constitutive heterochromatin (H3K9me3), facultative heterochromatin (H3K27me3), active enhancers and promoters (H3K27ac), and promoters at active genes (H3K4me3)^69,70^. We plotted signals for each of these PTMs via profile plots at all well-positioned nucleosomes (identified using MNase-seq inputs in each cell line), +1 nucleosomes (i.e., the first nucleosome downstream from a TSS), and nucleosomes bound by either PAX::FOXO1 fusion protein (**Figure 5A,B**). As expected given the relatively higher turnover rates, +1 nucleosomes had lower genomic signals for H3K9me3 and H3K27me3, but substantially higher genomic signals for H3K4me3 and H3K27ac^67^. Using these data and the average PTM signal at all well-positioned nucleosomes as a basis for comparison, nucleosomes bound by PAX3::FOXO1 have a relatively higher H3K9me3 and notably higher H3K27me3 signal (**Figure 5A**). In contrast, PAX7::FOXO1-bound nucleosomes had a higher signal for H3K4me3 and H3K27ac relative to those background nucleosomes (**Figure 5B**). This may be reflective of PAX7::FOXO1’s more prominent role at promoters with relatively higher basal activation states (**Figures 1G,H, 4, Supplemental Figure S3**).

**Figure 5.**
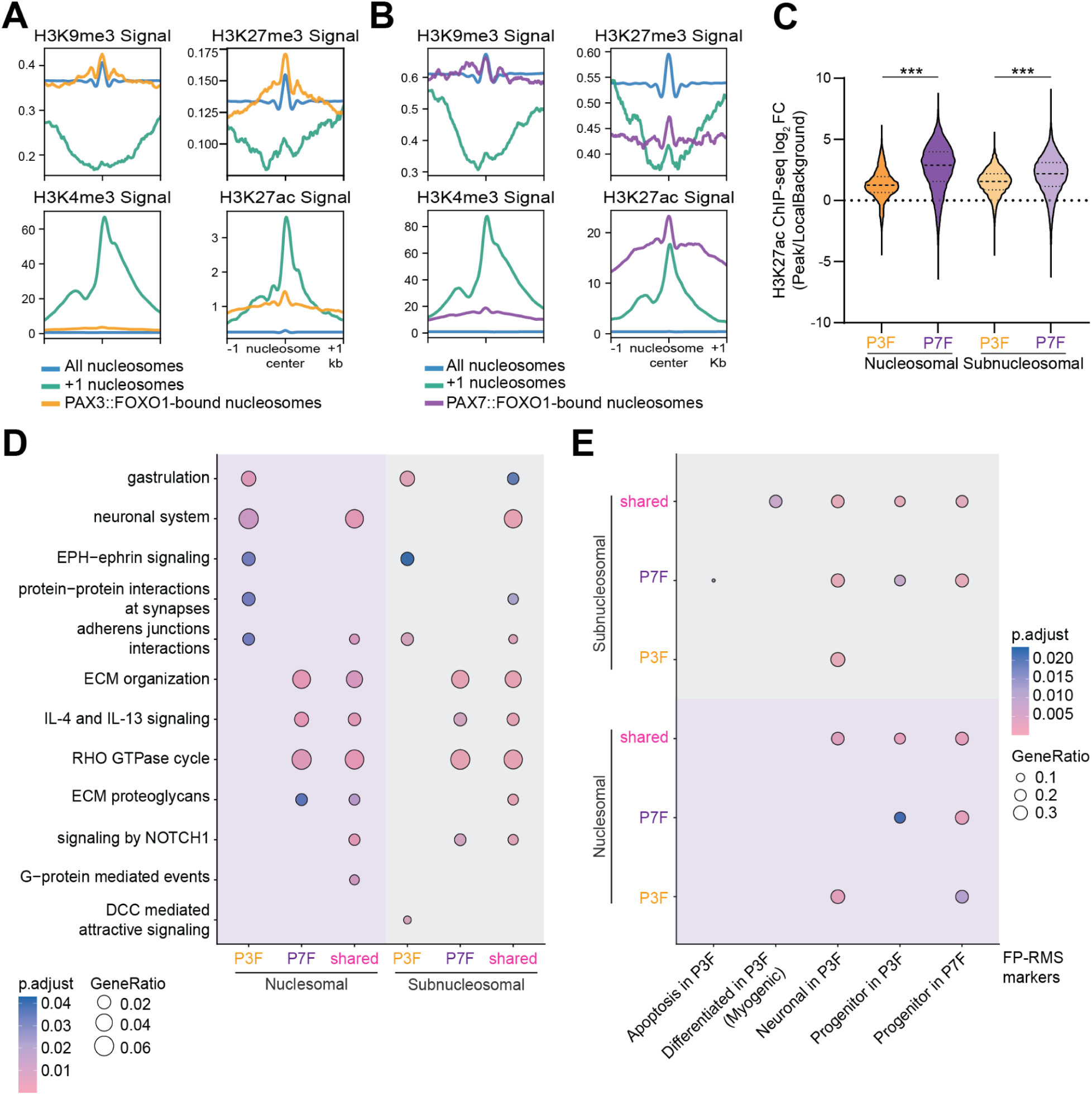
PAX7::FOXO1 has a higher association with active chromatin and distinct gene targeting preferences when binding nucleosomal chromatin in comparison to PAX3::FOXO1. (A) Profile plots comparing signal of histone post-translational modifications (PTMs) including H3K9me3, H3K27me3, H3K27ac, and H3K4me3 in RH4 cells at all well-positioned nucleosomes, those that are at the +1 position immediately downstream of a TSS, and those that are bound by PAX3::FOXO1 (P3F). (B) Profile plots comparing signal of histone PTMs (including H3K9me3, H3K27me3, H3K27ac, and H3K4me3) in CW9019 cells at all well-positioned nucleosomes, those that are at the +1 position immediately downstream of a TSS, and those that are bound by PAX7::FOXO1 (P7F). (C) Average quantified H3K27ac ChIP-seq signal between replicates at P3F MNase XChIP and P7F MNase XChIP binding sites in their respective RMS cell models. Dark dashed lines (---) represent means and thin dotted lines (···) represent quartiles. A Kruskal-Wallis test with a Dunn’s correction for multiple comparisons was used to determine statistical significance. (***) indicates *p* < 0.001. (D) Gene over-representation analysis (ORA) for reactome pathways on genes near differential P3F, P7F, and shared binding sites. Differential peaks were determined by DESeq2, FDR < 0.05 and nearby genes identified with GREAT (Genomic Regions Enrichment of Annotations Tool). (E) ORA to fusion-positive rhabdomyosarcoma markers, identified from previously reported single cell RNA-seq analysis (Danielli et al., 2024) for the same genes as in (D).

We calculated co-localization for these four PTMs (H3K9me3, H3K27me3, H3K27ac, H3K4me3). For each PAX::FOXO1 fusion, overlaps were identified between the called ChIP-seq peaks for each PTM and the three categories of nucleosomes from our MNase XChIP data (all well-positioned, all +1, and PAX::FOXO1 bound). We plotted the observed vs. expected ratios for each comparison, with results consistent with those seen in the profile plots (**Supplemental Figure S4A**). Nucleosomes bound by PAX3::FOXO1 had higher co-localization than expected for H3K9me3 and H3K27me3, while those bound by PAX7::FOXO1 had higher co-localization than expected for H3K4me3. We then calculated H3K27ac signal at nucleosomal and subnucleosomal PAX3::FOXO1 and PAX7::FOXO1 binding sites versus and observed that at both peak sets PAX7::FOXO1 had a greater relative enrichment for H3K27ac deposition both at its bound nucleosomes and accessible sites (**Figure 5C**). Importantly, H3K27ac is highly enriched at active promoters across the genome, along with H3K4me3. The higher observed over expected values for PAX7::FOXO1’s engagement with H3K27ac-positive nucleosomes could represent its having preferential engagement with active promoters, or a greater relative capacity to increase H3K27ac during its pioneering catalytic cycle. Similarly, we found that PAX7::FOXO1 sonication peaks had a greater overlap with H3K27ac than PAX3::FOXO1 (69% versus 45%) (**Supplemental Figure S4B**). Altogether, our analyses are consistent with PAX7::FOXO1 engaging promoter +1 nucleosomes as a stronger transcriptional activator, (see **Figure 1**), since in these patient-derived cell lines its binding often co-occurs and greater amounts of H3K27ac. While PAX3::FOXO1 may have distinct features due to its catalytic cycle of pioneering in its ability to engage farther toward the nucleosome dyad and relative greater affinity for nucleosomes within regions of repressive chromatin modifications.

### Differential binding of PAX3::FOXO1 and PAX7::FOXO1 fusions supports targeting of specific gene signatures

To understand the functional output of divergent modes of PAX3::FOXO1 and PAX7::FOXO1 chromatin engagement, we completed differential enrichment analysis on active chromatin marked by H3K27ac (ChIP-seq), between CW9019 (PAX7::FOXO1-positive) and RH4 (PAX3::FOXO1-positive) cells. CW9019 cells had 29,596 unique H3K27ac peaks and RH4 cells had 12,073, with 15,631 shared peaks between them (**Supplemental Figure S4C**). We investigated genes with differential H3K27ac levels at their promoters, given the high correlation between H3K27ac and gene expression^71–73^. In CW9019 cells, genes with greater H3K27ac at promoters were involved in the RHO GTPase cycle, β-catenin phosphorylation, and interleukin signaling. Meanwhile, RH4 cells had greater H3K27ac at myogenic, cellular senescence, and extracellular matrix genes (**Supplemental Figure S4D**). Sites with higher H3K27ac in CW9019 cells were highly associated with bZIP motifs, and RH4-enriched H3K27ac sites were highly associated with bHLH motifs (**Supplemental Figure S4E**). This finding may suggest different preferences in binding to the degenerate composite PD-HD motif, which includes the bZIP motif, for which PAX7::FOXO1 has a higher preference (**Figure 3D**). Alternatively, this motif analysis could also represent a difference in cooperativity with those transcription factor families, which will be a focus of future studies.

We also completed differential binding analysis between nucleosomal and subnucleosomal PAX3::FOXO1 binding sites and PAX7::FOXO1 binding sites from MNase XChIP in their respective cells. We identified potential gene targets of these binding sites with the Genomic Regions Enrichment of Annotations Tool (GREAT) for pathway analysis. Comparing nucleosomal binding revealed that PAX3::FOXO1 was better able to bind near neuronal and synapse-interacting genes, and PAX7::FOXO1 better bound near interleukin signaling, RHO GTPase cycling, and extracellular matrix genes (**Figure 5D**, cf., **Supplemental Figure S4D**). Subnucleosomal binding of PAX7::FOXO1 is associated with a similar gene set to the nucleosomal sites, while PAX3::FOXO1 subnucleosomal binding occurred near ephrin signaling genes. Next, we did pathway analysis for FP-RMS marker genes. Nucleosomal PAX3::FOXO1 engagement occurred near neuronal cell state markers, and PAX7::FOXO1 nucleosomal binding was more prevalent at progenitor cell state markers (**Figure 5E**). Subnucleosomal PAX7::FOXO1 peaks were, again, at progenitor cell state markers, and neuronal marker binding appears consistently across PAX::FOXO1 subnucleosomal peaks. Altogether, PAX3::FOXO1 nucleosomal binding supports further and stronger binding near neuronal pathways and markers, a capability that subnucleosomal peaks do not enhance. Therefore, the improved nucleosomal binding by PAX3::FOXO1 may provide a new avenue for PAX3::FOXO1 to access and potentially activate these transcriptional programs.

## DISCUSSION

PAX-fusion oncoproteins are known to drive a more aggressive and specific histological subtype of rhabdomyosarcoma. However, despite the similarities between PAX3::FOXO1 and PAX7::FOXO1, there are significantly different patient outcomes based on the driving fusion, with PAX3::FOXO1 associated with a worse prognosis. Here, we identify that these PAX-fusion oncoproteins rapidly drive initial transcriptional programs in zebrafish with key similarities but distinct functional outputs (**Figure 1**). We were able to determine key differences in the chromatin binding by endogenous PAX3::FOXO1 and PAX7::FOXO1 fusions through comparative analysis of a modified MNase XChIP protocol to capture fusion/nucleosome engagement events as well as events with accessible chromatin. Therefore, these divergent clinical outcomes, and distinct immediate-early gene activation programs, may be driven not by specific transcriptional programs but through distinct pioneering and chromatin recognition functions of PAX3::FOXO1 and PAX7::FOXO1.

Our modified MNase XChIP protocol enabled us to decipher the differential engagement of exposed DNA, and nucleosomal DNA. While each PAX-fusion oncoprotein is capable of binding DNA in either context, they appear to do so through distinct modalities (**Figure 2D**). At nucleosomes, PAX3::FOXO1 preferred to bind at composite PD-HD motifs (RTGACTAAT), which can occur across the body of the nucleosome and at entry/exit sites. Meanwhile, PAX7::FOXO1 preferred to bind to a degenerate composite motif (RTGASTAAT) strictly at nucleosomal entry/exit sites (**Figures 3C,D, 4**). A study on wild-type PAX7, which has the same DNA binding domains as the fusion oncoprotein, uncovered enrichment of this degenerate motif specific to its function in melanotrope cells^23^. PAX3::FOXO1 binding sites also contain the degenerate composite motif, but not to the same extent as PAX7::FOXO1. Both PAX::FOXO1 fusion proteins in RMS show notable binding to a longer composite-related motif compared to binding at partial motifs (**Figure 3C**). This binding propensity parallels a proposed model by which some pioneer factors overcome the energetic barriers presented by nucleosomes^35^. However, composite motif identification was not the predominant mode when PAX3::FOXO1 was binding inaccessible chromatin in zebrafish or upon its sampling of sites in inducible myoblasts^28,43^. In those studies, we observed binding to homeobox and bZIP motifs, respectively. It remains unclear what underlying secondary layer of logic and factors influence PAX::FOXO1 fusion proteins or other pioneer factors to shift their mode of interaction for closed chromatin between experimental conditions or biological contexts. Possibilities could include endogenous nucleosome positioning sequences^74^, nucleosome turnover rates^67,75^, cooperativity^68,76^, and 3D architectural constraints^27,77^.

We found that PAX7::FOXO1 had a lower proportion of nucleosomal binding sites compared to PAX3::FOXO1 (**Figure 3E-G**). Additionally, PAX3::FOXO1 displayed an increased overlap with repressive histone marks, H3K27me3 and H3K9me3 (**Figure 5A,B, Supplemental Figure S4A**). There are several potential explanations for these differential localization patterns, including (1) PAX3::FOXO1 may have a better capacity to invade closed chromatin, or (2) PAX3::FOXO1 and PAX7::FOXO1 fusions have similar binding capabilities, but PAX7::FOXO1 activates a greater majority of targets residing in repressed chromatin^31^. However, we find the second possible explanation to be less likely given that a greater fraction of PAX7::FOXO1 gene targets have a higher basal activation state (**Figure 1G,H**). The distinct chromatin regulatory preferences of PAX3::FOXO1 and PAX7::FOXO1 could influence when or whether specific transcriptional programs are activated, such as metastatic or therapeutic resistance programs in a particular biological context. Electrophoretic mobility shift assays or domain swap experiments could further quantify relative binding motif affinities and assess how each domain individually contributes to chromatin invasion and gene activation mechanisms. For example, since wild-type PAX7 has a greater affinity to homeobox motifs than wild-type PAX3, the positioning of the homeodomain could be a particular driver of differential nucleosome invasion^18^. It also remains possible that binding to closed chromatin is influenced by the conserved C-terminus of FOXO1, which has demonstrated direct protein-protein contacts with the H3-H4 tetrameric core of nucleosomes and capacity for invading H1-compacted heterochromatin^78^. How the combinatorial complexity of DNA-binding domains and activator domains tunes the thermodynamics of chromatin binding will be a focus of future work, including investigations of how the C-terminal domains function in concert with the PD-HD domains of the fusions. In particular, such efforts could determine if PAX7::FOXO1’s preference for binding degenerate motifs may be due to direct structural differences from PAX3::FOXO1 or higher order chromatin structure. Together, our study illuminates and measures fusion/nucleosome binding in RMS cells, defines chromatin context including initial nucleosome positioning information in RMS, and reveals key distinctions in fusion targeting which are in alignment with observed clinical and in vivo trends.

## Limitations of the study

We examined PAX3::FOXO1 and PAX7::FOXO1 in patient-derived cell lines, which are a steady-state system that is already transformed. Therefore, we cannot decouple which binding sites were already accessible in the background environment versus those activated by PAX::FOXO1 fusions in our system. Furthermore, there could be different binding modes between sites that were activated or bound by PAX::FOXO1 fusions, but not activated, that our system has not captured. These differences could represent alternative mechanisms for tumor initiation and maintenance. Understanding the functions of PAX::FOXO1 fusion oncoproteins across developmental contexts is especially critical, because it remains unknown when the somatic fusion oncogene is acquired and the latency to tumor presentation. The retained *PAX3/PAX7* promoter could drive different regulatory timing of expression, potentially and likely providing divergent cells of origin for these oncoproteins. The application of MNase XChIP to dynamic systems of inducible expression or degradation across developmental contexts could further elucidate additional mechanisms of binding and connect specific chromatin events to transcriptional consequences.

## STAR*METHODS

### RESOURCE AVAILABILITY

Further information and requests for resources and reagents should be directed to and will be fulfilled by the lead contact, Benjamin Z. Stanton.

### MATERIALS AVAILABILITY

This study did not generate new, unique reagents.

### DATA AND CODE AVAILABILITY

- All data reported in this paper will be shared by the lead contact upon request.
- Sequencing data will be deposited on GEO following revisions and acceptance of the manuscript.
- Original codes to analyze the data from this paper will be posted on GitHub following revisions and acceptance of the manuscript.
- Any additional information or raw data required to reanalyze the data reported in this paper is available from the lead contact upon request.

### EXPERIMENTAL MODEL AND STUDY DESIGN

#### Zebrafish Husbandry

Zebrafish (*Danio rerio)* were maintained in an AAALAC-accredited, USDA-registered, OLAW-assured, and Guide for the Care and Use of Laboratory Animals compliant facility at Nationwide Children’s Hospital, as previously described^28^. Adult zebrafish are housed in tanks with a density of 5–12 fish per liter on a recirculating system in 28°C carbon-filtered water. Fish from 5 to 30 days post-fertilization were fed live rotifers three times a day and a commercial pelleted diet twice daily, after 30 days post-fertilization. Micro-injections in zebrafish embryos were performed on male and female zebrafish. WIK zebrafish were used as wild-type and were obtained from the Zebrafish International Resource Center (ZIRC; https://zebrafish.org/). Research procedures were approved by the IACUC at The Abigail Wexner Research Institute at Nationwide Children’s Hospital.

#### Cell Culture

Rhabdomyosarcoma cell lines RH4 (PAX3::FOXO1-positive rhabdomyosarcoma) and SMS-CTR (fusion-negative rhabdomyosarcoma) were gifts from Dr. Peter Houghton at UT Health San Antoni, and CW9019 (PAX7::FOXO1-positive rhabdomyosarcoma) were gifts from Beat Schäfer at University Children’s Hospital Zurich. All cells were cultured in DMEM supplemented with Glutamax, FBS (10% v/v), and penicillin-streptomycin.

## METHOD DETAILS

### Plasmid Cloning and mRNA Injection Construct Generation

The human PAX7::FOXO1 coding sequence was synthesized from the coding sequence previously reported by Davis et al^13^. Primers for plasmid cloning are listed below:

- FWD: GAGGTTAATTAAGCCGCCACCATGGCGGCCCTTCCCGGC
- REV: GATCGGCGCGCCGCCTGACACCCAGCTATG

*PacI* and *AscI* restriction digest sites were added to the PCR for cloning into pCS2+MCS-P2A-sfGFP with T4 DNA ligase (NEB, M0202S) in a 3:1 insert to backbone ratio. The backbone plasmid was a kind gift from Jason Berman (Addgene plasmid #74668). Ligated plasmids were transformed into DH5α cells (Fisher Scientific, FEREC0111), linearized with *NotI*, for in vitro mRNA transcription with the mMESSAGE mMACHINE SP6 Transcription Kit (ThermoFisher Scientific, AM1340). PAX3::FOXO1 (Plamsid# 240098) and CNTL (Plasmid# 74668) mRNA constructs were generated as previously reported^28^. PAX7::FOXO1 plasmid will be deposited to Addgene.

### Zebrafish Embryo Injection

Adult wild-type WIK zebrafish were in-crossed. Fertilized eggs were injected in the yolk at the single-cell stage with either 100 ng/μL of PAX7::FOXO1–2A-sfGFP, equimolar of PAX3::FOXO1–2A-sfGFP, or equal molarity of the CNTL (-2A-sfGFP backbone), with a drop size diameter of 0.15 mm. Injection mixes consisted of mRNA and 0.05% phenol red in 3X Danieau’s buffer (52.2 mM NaCl, 0.63 mM KCl, 0.36 mM MgSO_4_·7H_2_O, 0.54 mM Ca(NO_3_)_2_·7H_2_O, 4.5 mM HEPES). The embryos were incubated in 1X E3 buffer (5 mM NaCl, 0.17 mM KCl, 0.33 mM CaCl_2_, 0.33 mM MgSO_4_) at 32°C until they reached the 6-hour post-fertilization shield stage according to Kimmel et al.^79^. Embryonic survival was completed by counting the initial number of injected embryos and the number of dead embryos at each labeled developmental time point.

### Zebrafish Western Blotting

Embryos were collected at the 6-hour post fertilization timepoint. Embryos were dechorionated with pronase, deyolked (55 mM NaCl, 1.8 mM KCl, 1.25 mM NaHCO_3_), and washed with 0.5X Danieau’s buffer (29 mM NaCl, 0.35 mM KCl, 0.2 mM MgSO_4_·4H_2_O, 0.3 mM Ca(NO_3_)_2_·4H_2_O, 2.5 mM HEPES). Embryos were snap-frozen with dry ice and stored at −80°C. Protein lysates were generated by adding 2 μL of 2X Laemmli Buffer (Bio-Rad, 1610737) per embryo and heating samples for 5 min at 95°C. Equal volume and embryo number were loaded. Protein was transferred to a 0.2 μm PVDF membrane (Bio-Rad, 1620177), blocked with Casein and 0.05% Tween 20 (Fisher Scientific, PI37528), and incubated with primary antibodies. αFOXO1 (Cell Signaling, C29H4) was used for PAX3/7::FOXO1 and αTUBULIN (Cell Signaling, 3873S) for Tubulin. Secondary antibodies of HRP-α-rabbit (Bio-Rad, 1721019) and HRP-α-mouse (Bio-Rad, 1706516), respectively. Images were taken on a ChemiDoc Go Imaging System (Bio-Rad, 12018025) with the SuperSignal West Atto Ultimate Sensitivity Chemiluminescent Substrate (Fisher Scientific, PIA38554).

### RNA-sequencing (RNA-seq)

Embryos were injected and collected as described above. A minimum of 20 embryos were injected, and replicates were generated across multiple injection days. RNA was isolated with a QIAGEN RNeasy Mini kit with on-column DNase digestion (74104, QIAGEN). NEBNext Ultra II Directional RNA Library Prep Kit for Illumina (E7760, New England Biolabs) and a polyA enrichment step (E7490, New England Biolabs) were used for library preparation. RNA samples were run on a NovaSeq SP with 150bp paired-end reads at the Nationwide Children’s Hospital Institute for Genomic Medicine (IGM). RNA-seq alignment, gene counts, and differential expression analysis were completed with the nf-core/rnaseq (10.5281/zenodo.1400710) and nf-core/differential abundance pipelines (10.5281/zenodo.7568000)^80^. Pathway analysis was completed with clusterProfiler^81^ and MSigDB database^82^. Other plots were generated with ggplot2^83^ and ComplexHeatmap^84^. Additional statistical analysis was completed in GraphPad Prism 10 and R (version 4.3.0), and additional experimental information is included in the respective figure legends.

### Cellular Fractionation and Immunoblotting

Sonication-based fraction was completed as previously described^43^. Cells were fractionated into cytoplasmic, nuclear, soluble chromatin, and insoluble chromatin pellets. For salt-based fractionation, cells were treated with an increasing gradient of salt concentrations according to ref.^44^. Cytoplasmic, freely diffusing nuclear, euchromatin-associated, and heterochromatin-associated fractions were collected. Cell fractions were quantified with the Pierce Rapid Gold BCA Protein Assay Kit (ThermoFisher Scientific Cat# A55860). Samples were resolved by SDS-PAGE on a NuPAGE 4-12% gradient Bis-Tris gel. Proteins were transferred overnight at 4°C with a constant voltage of 24V or for one hour at 100V onto nitrocellulose membranes. Membranes were blocked at room temperature in 5% w/v milk solution in TBST (0.1% v/v Tween-20) for 1 hour before incubating for 2 hours at room temperature with primary antibodies detecting: BAF155 (Cell Signaling Technology Cat# 11956, RRID:AB_2797776), FOXO1 (Cell Signaling Technology Cat# 72874, RRID:AB_2799829), HP1(α/β) (Cell Signaling Technology Cat# 2623, RRID:AB_2070981) or TBP (Cell Signaling Technology Cat# 8515, RRID:AB_10949159). Membranes were incubated for a further 1 hour with goat anti-rabbit IgG (H+L) secondary antibody, HRP (Thermo Fisher Scientific Cat# 31460, RRID:AB_228341). Membranes were imaged with the ChemiDoc Go Imaging System (Bio-Rad, Cat# 12018025) after incubation with SuperSignal West Pico PLUS Chemiluminescent Substrate (Thermo Fisher Scientific Cat# 34577).

### Modified MNase XChIP

Cells were cultured as described above, and 6×10^6^ cells were collected. Cells were dissociated in microcentrifuge tubes and centrifuged at 300g for 5 minutes at room temperature (RT). The supernatant was removed, and the pelleted cells were re-suspended in PBS and fixed in 1% formaldehyde for 10 minutes at RT. Cells were inverted every 2-3 minutes to ensure thorough and even fixation. Fixation was quenched by the addition of a glycine solution to a final concentration of 125 mM. The samples were incubated on ice for 5 minutes and inverted to ensure thorough quenching. Fixed cells were pelleted again via centrifugation at 4°C, 1,200g for 5 minutes. The supernatant was removed, the cells were washed in PBS containing proteinase inhibitors, and stored at −80°C following snap-freezing.

Fixed cells were thawed slowly on ice and resuspended in PBS supplemented with proteinase inhibitors. Cells were centrifuged at 4°C, 1,200g for 3 minutes. The supernatant was discarded, and the cells were resuspended in 100 μL lysis buffer (1% SDS, 10 mM EDTA, 50 mM Tris, pH 8 in ddH_2_O) supplemented with proteinase inhibitor and incubated on ice for 10 minutes. 1.66 μL 2M CaCl_2_ and proteinase inhibitor were freshly added to 1 mL ChIP Dilution Buffer (1% Triton-X, 2mM EDTA, 150 mM NaCl, 20 mM Tris in ddH_2_O). 900 μL of complete ChIP Dilution Buffer was added to lysed cells for a total volume of ∼1 mL. Diluted cells were mixed by gentle pipetting and incubated at 37°C for 2 minutes. 3.33 μL of 2x10^6^ U/mL MNase was diluted separately in 40 μL ChIP Dilution buffer before being added to the cell lysate and mixed well by inverting 4-5 times. The digestion reaction was incubated for 15 minutes at 37°C. The digestion reaction was stopped rapidly by the addition of pre-mixed 30 μL EDTA (0.5 M) and 60 μL EGTA (0.5 M) before being placed directly on ice. The digested cells were sonicated gently for 30 seconds to aid in the solubilization of chromatin. The digested and solubilized chromatin was centrifuged at 16,000g for 2 minutes. The supernatant was removed and kept as 40 μL aliquots for use as input chromatin. To decrosslink the input chromatin, 186 μL H2O, 3 μL Tris (1 M), 6 μL EDTA (0.5 M), 12 μL EGTA (0.5 M), 18 μL NaCl (5 M), 30 μL 10% SDS, and 5 μL proteinase-K were added and incubated overnight at 65°C. The remaining supernatant was divided into two 400 μL aliquots for immunoprecipitation. 5 μL of anti-FOXO1 antibody (Cell Signaling Technology Cat# 72874, RRID:AB_2799829) was added and incubated overnight at 4°C with end-over-end rotation. 40 μL of Dynabeads were resuspended in ChIP Dilution buffer and added to the antibody-incubated chromatin. The mixture was incubated for an additional 3 hours at 4°C with end-over-end rotation. Beads were quickly washed at room temperature with ice-cold buffers in the following order: 2x washes TSE (0.1% SDS, 1% Triton-X, 2 mM EDTA, 20 mM Tris pH 8, 150 mM NaCl in ddH_2_O); 2x washes Buffer 3 (0.25 mM LiCl, 1% NP-40, 1% DOC, 1 mM EDTA, 10 mM Tris pH 8 in ddH_2_O); 2x washes TE (pH 8). For each wash, the beads were isolated by magnet and resuspended by end-over-end rotation. Isolated beads were resuspended in 100 μL TE (pH 8), 2.5 μL 10% SDS, and 5 μL proteinase-K following the final wash. Chromatin was decrosslinked from beads by incubating overnight at 65°C. Beads were removed via magnet, and decrosslinked input and IP chromatin were separately purified via QIAQuick spin column and eluted in 36 μL pre-warmed elution buffer (EB, Qiagen Cat# 19086).

### Modified MNase XChIP Library Preparation

MNase XChIP libraries were prepared as previously reported^41^ with some modifications to ensure isolation and preservation of nucleosomal and TF-footprint-sized DNA fragments. Samples used as starting material: 2 µL of input or 10 µL of immunoprecipitated DNA diluted in 83 µL or 75 µL ddH_2_O, respectively. 10 µL of 10X NEBNext® End Repair Reaction Buffer and 5 µL NEBNext® End Repair Enzyme Mix were added to each sample (for 100 µL total per sample) and incubated for 30 minutes at 20°C before being purified via QIAQuick spin column and eluted in 32 µL EB. A-tailing was completed with the addition of 10 µL 1 mM dATP, 5 µL NEBuffer 2, and 3µL Klenow fragment (3’-5’ exo-) to the eluted end-repaired DNA (for 50 µL per sample), incubated for 30 minutes at 37°C, and purified via QIAQuick spin column and eluted in 22 µL EB. 3 µL 10X T4 DNA Ligase Buffer, 3 µL T4 DNA Ligase, and 2 µL PE adapter mix (pre-annealed at 15 µM concentration) were added to the 22 µL of A-tailed DNA and mixed gently by pipette before allowing to incubate for 30 minutes at RT. 30 µL of the adapter-ligated DNA mixture was then added to 54 µL of AMPure XP magnetic beads (1 to 1.8 DNA to bead ratio) and allowed to incubate for 5 minutes at RT. Beads were isolated and washed twice with fresh 80% ethanol. Beads were air-dried for 10 minutes before being gently dissolved in 25 µL ddH_2_O. 23 µL of the dissolved DNA was mixed with unique combinations of 1 µL of Illumina i5 index adapter, 1 µL of Illumina i7 index adapter, and 25µL of NEB Phusion™ High-Fidelity PCR Master Mix (for 50 µL total per sample). The PCR amplification used only 25 µL of this reaction (the remaining volume can be stored at -20°C until needed). Samples were amplified for 12-16 cycles (each cycle consisting of: 98°C for 10 seconds, 65°C for 30 seconds, 72°C for 30 seconds), with cycle number minimized to avoid PCR duplicates while ensuring clearly discernible bimodal distributions of TF-footprint/subnucleosomal and nucleosomal-sized fragment lengths.

### In Silico Sorting of Modified MNase XChIP Sequencing

MNase XChIP fastq files were trimmed using Cutadapt (v.3.4)^85^ and FastQC(v0.12.1) before being sorted using Cutadapt. Nucleosomal reads were isolated using a minimum cutoff “-m 130” and subnucleosomal reads using a maximum cutoff “-M 86” and option “--pair-filter=any” to require both reads in a pair meet these requirements. These sorted fastq files were then carried forward for downstream analysis.

### ChIP Sequencing

ChIP-seq experiments were done as previously described^27,43^. Immunoprecipitations used the following antibodies: FOXO1 (Cell Signaling, Cat#72874), H3K27ac (Active Motif, Cat#39133), H3K4me3 (Active Motif, Cat#61379), H3K27me3 (Active Motif, Cat#39155), and H3K9me3 (Active Motif, Cat#39062). H3K9me3 and H3K27ac ChIP-seq in RH4 cells were meta-analyzed from Wang et al., 2023 and Sunkel et al., 2021.

### ChIP-seq and MNase XChIP-seq Data Analysis

ChIP-seq and sorted MNase XChIPs were analyzed using nf-core’s chipseq pipeline (v.2.1.0)^80^, with the Bowtie2 aligner^86^ to the hg38 genome. Peaks were called using 0.005 for the “--macs_pvalue” parameter. Consensus peaks were called using a minimum overlap of 2 (for sonication ChIPs) or 3 (for MNase XChIPs) replicates. Bigwigs were generated using a custom PerCell implementation^41^ of MACS2’s bdgcmp^87^ to enable incorporation of input into signal tracks. Well-positioned nucleosomes were called using the nucleR software^88^ with width and height cutoffs of 0.9 for parameters “--wthresh” and “--hthresh.” De novo and known motif enrichments were identified using HOMER^63^. Signal tracks were visualized using Trackplot^89^. UpSet plots^90^ were generated using UpSetR software (https://doi.org/10.1093/bioinformatics/btx364) using intersections calculated using HOMER’s mergePeaks command with parameter “-d given.” Observed vs. expected ratios for peak set overlaps were determined using the same command. Profile plots and heat maps were generated using computeMatrix and plotHeatmap from deepTools (v. 3.5.6). Matrices for the heat maps were sorted in decreasing order based on average signal in the central 150 bp region. Occupancy plots were created using plot2DO (Bereti and Chereji, 2020). Putative binding motifs were identified within peaks using HOMER’s scanMotifGenomeWide and intersecting with peaks via BEDTools intersect (v.2.30.0)^91^.

## ACKNOWLEDGMENTS

We thank all members of Stanton and Kendall Labs for helpful discussions and comments. We are grateful to the Nationwide Children’s Hospital (NCH) Animal Resources Core for their exceptional zebrafish husbandry, especially the Zebrafish Facility team, Dr. Carmen Arsuaga, Logan Fehrenbach, Logan Bern, and Jacob Al-Armanazi. We thank Dr. Meng Wang for helpful discussions related to this work. We are grateful to the Steve and Cindy Rasmussen Institute for Genomic Medicine Genomics Services Laboratory for assistance with sequencing, and the High-Performance Computing Group for assistance maintaining and using the NCH cluster.

B.Z.S. is grateful for support from the National Cancer Institute (U01CA298763), National Institute of General Medical Sciences (R01GM144601), National Heart Lung and Blood Institute (R01HL166520), American Cancer Society (RSG-23-1021178-01-DMC), St. Baldrick’s Foundation (Childhood Cancer Research Grant), and Nationwide Children’s Hospital. G.C.K. is grateful for support from an NIH/National Cancer Institute R01 CA272872 grant, an Alex’s Lemonade Stand Foundation “A” Award, a CancerFree Kids New Idea Award, a DOD Cancer Idea HT9425-25-1-0356 Award, and Nationwide Children’s Hospital. J.K. and R.A.H are supported by a T32 CA269052 Training Program in Basic and Translational Pediatric Oncology Research predoctoral fellowship. C.T. is thankful for support from a CancerFree Kids New Idea Award. The Institute for Genomic Medicine is funded by the Nationwide Foundation Pediatric Innovation Fund and the Ohio State University Comprehensive Cancer Center grant P30 CA016058. The funders had no role in study design, data collection and analysis, decision to publish, or preparation of the manuscript. The content is solely the responsibility of the authors and does not necessarily represent the official views of the NIH.

## AUTHOR CONTRIBUTIONS

Conceptualization, A.T., J.K., G.C.K., and B.Z.S. Methodology, A.T., J.K., A.M.V., B.D.S., R.A.H., B.Z.S., and C.K. Modified MNase XChIP methodology was developed by A.T. and B.Z.S. Formal analysis, A.T., J.K, M.W., and C.T. Data curation and visualization, A.T. and J.K. Supervision, Project administration, and funding acquisition, G.C.K. and B.Z.S. Writing – original draft, A.T., J.K., G.C.K, and B.Z.S. Writing – review and editing, A.T., J.K., B.Z.S., C.T., G.C.K, and R.A.H. All authors reviewed and edited the final manuscript.

---

**Supplemental Figure S1.**
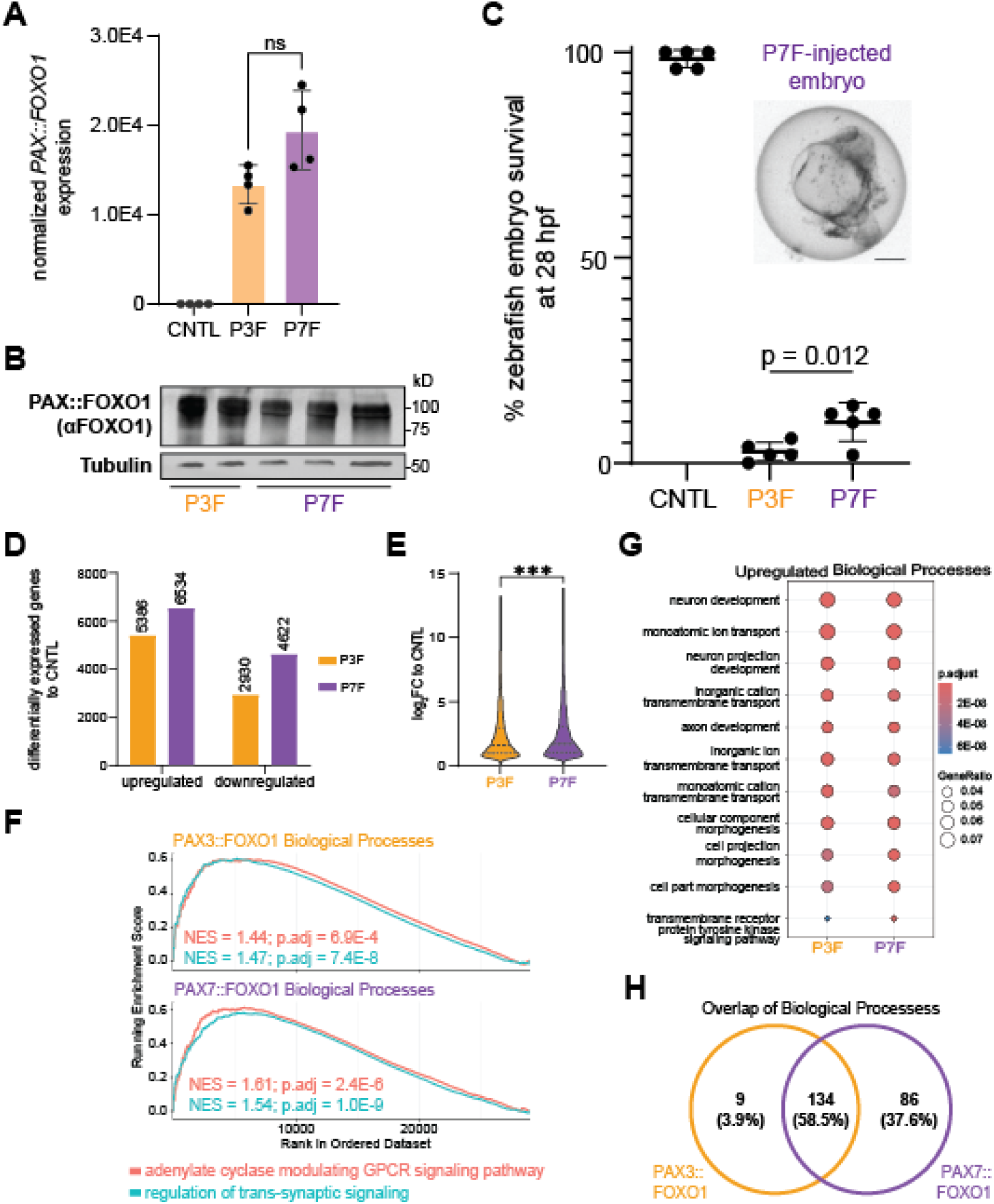
PAX7::FOXO1 drives stronger gene activation in zebrafish, and results in reduced lethality compared to PAX3::FOXO1. (A) Normalized fusion gene mRNA expression in control (CNTL), PAX3::FOXO1 (P3F), and PAX7::FOXO1 (P7F) in the respective zebrafish embryos. Each point represents gene expression in RNA-seq replicates. A Mann-Whitney test was used to determine statistical significance, and ns indicates not statistically significant. (B) Representative western blot of PAX3::FOXO1 expression and PAX7::FOXO1 expression in respective injected zebrafish embryos. Each lane is a technical replicate. (C) Survival of embryos injected with CNTL, P3F, or P7F mRNA. Each point represents survival of 50 embryos from one independent injection day. An ordinary one-way ANOVA with Tukey’s multiple comparisons test determined statistical significance. The upper right panel is a representative image of P7F embryos at 28 hours post-fertilization (hpf). Scale bar = 250 µm. (D) Plot of differentially expressed genes (absolute log_2_ fold change ≥ 1, p.adjust ≤ 0.05) between P3F or P7F versus CNTL. (E) Violin plot of log_2_FC versus CNTL of upregulated genes in E. Dark dashed lines (---) represent means and thin dotted lines (**···**) represent quartiles. A Mann-Whitney test was used to determine statistical significance. (***) indicates *p* < 0.001. (F) GSEA to adenylate cyclase modulating GPCR signaling pathway and regulation of trans-synaptic signaling biological processes for P3F or P7F versus CNTL. (G) Gene over-representation analysis of top upregulated genes between P3F or P7F versus CNTL. (H) Overlap of upregulated biological processes between P3F or P7F versus CNTL, pathway *p*-value cutoff < 0.01.

**Supplemental Figure S2.**
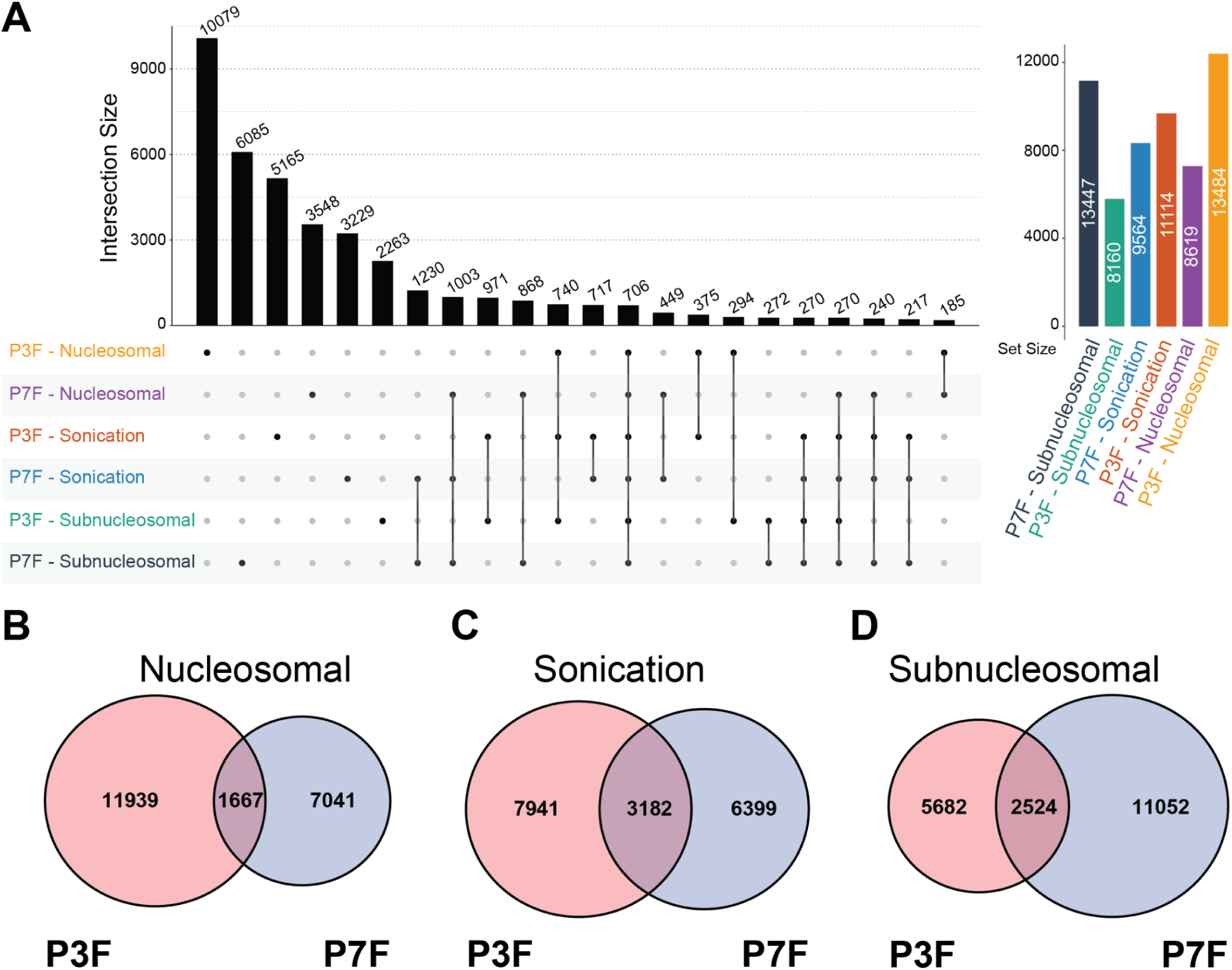
Nucleosomal (MNase XChIP targets), subnucleosomal (MNase XChIP targets), and sonication (ChIP-seq targets) reveal many unique binding sites between PAX3::FOXO1 and PAX7::FOXO1 fusion proteins. (A) UpSet plot displaying the number of unique and shared peaks across nucleosomal and subnucleosomal MNase XChIPs and sonication ChIPs for PAX3::FOXO1 (P3F) and PAX7::FOXO1 (P7F). For readability, only intersections with >150 peaks are shown. (B) Venn diagram overlap of nucleosomal P3F and P7F peaks. (D) Venn diagram overlap of sonication P3F and P7F peaks. (D) Venn diagram overlap of subnucleosomal P3F and P7F peaks.

**Supplemental Figure S3.**
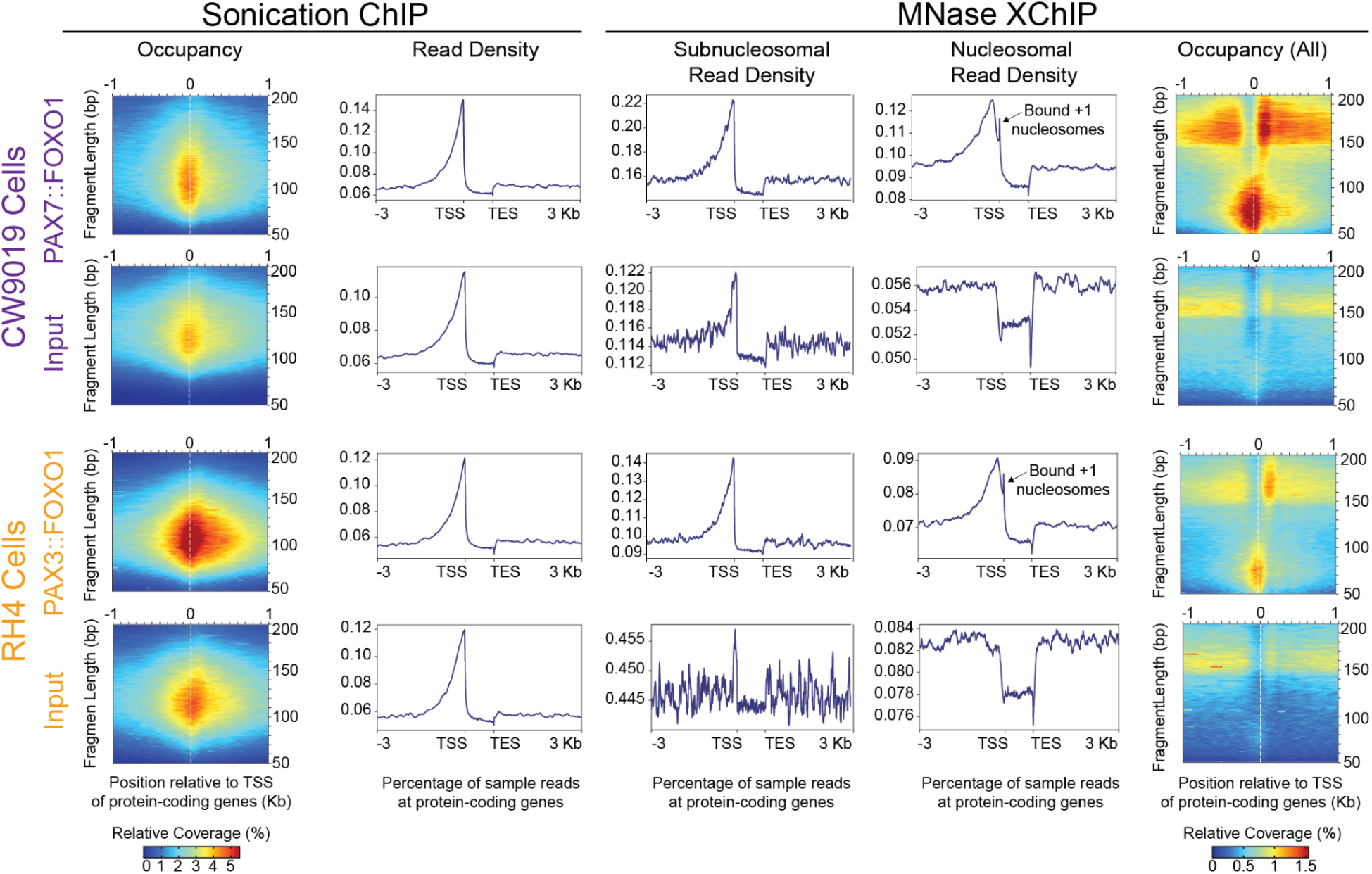
PAX3::FOXO1 and PAX7::FOXO1 bind at transcriptional start sites. Sonication ChIP for PAX7::FOXO1 (top, left), Sonication ChIP for PAX3::FOXO1 (bottom, left). Modified MNase XChIP for PAX7::FOXO1 (top, right), modified MNase XChIP for PAX3::FOXO1 (bottom, right), plot2DO occupancy plots showing signal +/- 1 kilobase (Kb) from gene transcriptional start sites (TSSs), including PAX7::FOXO1, PAX3::FOXO1, and their respective input samples. Genic regions are centered on TSSs on the x-axis with sequencing reads separated by fragment length on the y-axis and relative read coverage shown on the heatmap. Read coverage is the average of all replicates. Read density as percentages of sonication ChIP (left), subnucleosomal modified MNase XChIP (center, middle), and nucleosomal modified MNase XChIP (center, right). Regions extend 3 Kb upstream from TSS and 3 Kb downstream from transcriptional end sites (TES) of protein-coding genes from a representative replicate. Peaks in nucleosomal PAX::FOXO1 immunoprecipitations corresponding to bound +1 nucleosomes are indicated.

**Supplemental Figure S4.**
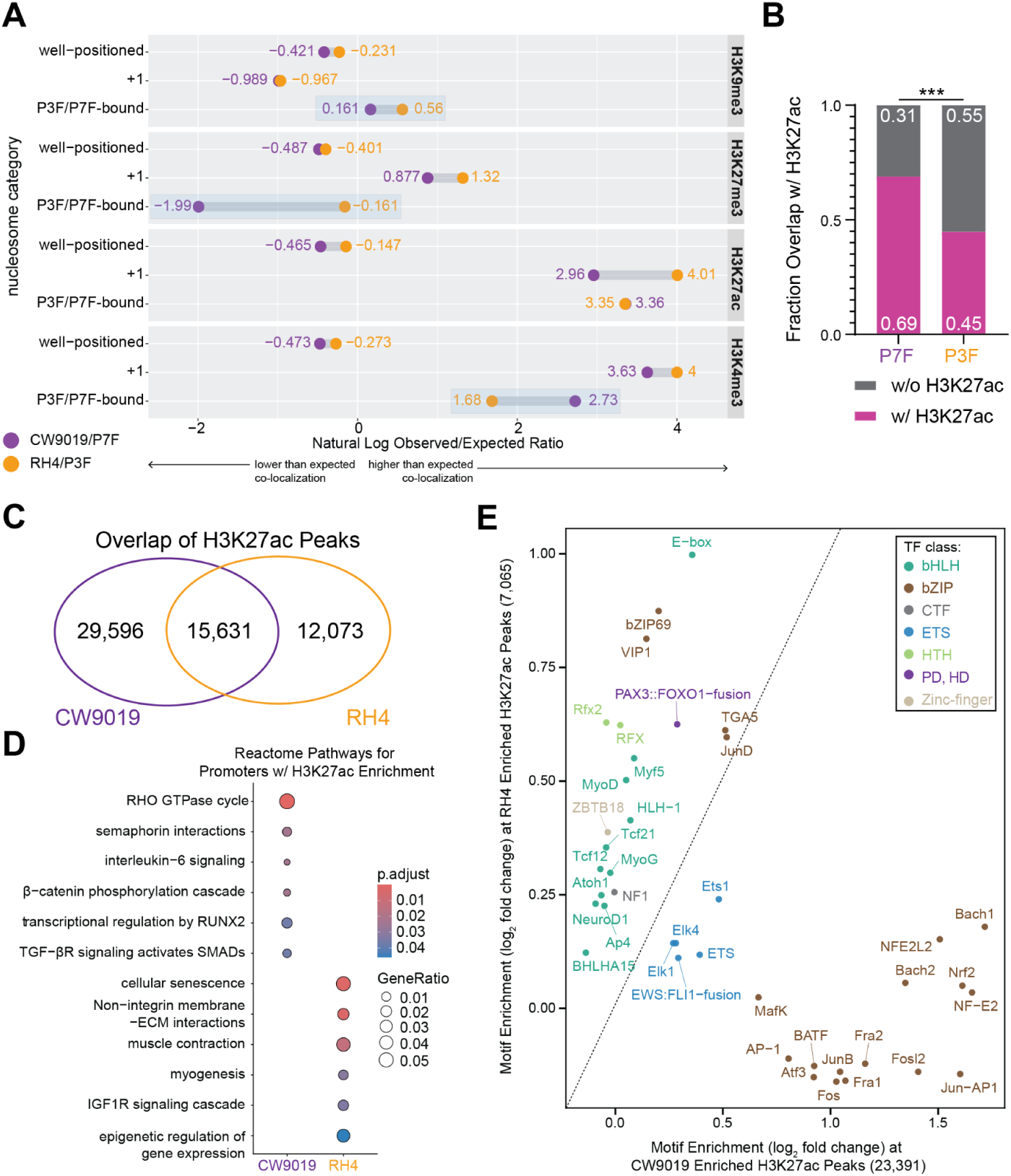
CW9019 and RH4 cells have distinguishing characteristics within their H3K27ac landscapes. (A) Observed/expected ratios for the co-occurrence of the indicated category of nucleosomes (MNase XChIP input data) with meta-analyses of H3K9me3, H3K27me3, H3K27ac, or H3K4me3 (ChIP-seq data) overlapping with called nucleosome-bound fusion target sites in RH4 cells (PAX3::FOXO1 MNase XChIP) or CW9019 cells (PAX7::FOXO1 MNase XChIP). Ratios for both cell lines are presented on the same row, with CW9019 shown in purple and RH4 in orange. Key contrasts are highlighted when PAX7::FOXO1 (P7F)-bound nucleosomes and PAX3::FOXO1 (P3F)-bound nucleosomes notably differ. (B) Overlap between P7F and P3F sonication ChIP-seq peaks with H3K27ac in CW9019 and RH4 cells, respectively. A Fisher’s exact test was used to determine statistical significance. (***) indicates *p* < 0.001. (C) Overlap of consensus H3K27ac ChIP-seq peaks between CW9019 and RH4 cells. (D) Gene over-representation analysis of genes with significantly differential H3K27ac levels at their promoters between CW9019 and RH4 cells, DESeq2, FDR< 0.01. (E) Known HOMER motif analysis of top 20 most significant motifs at all peaks with differential H3K27ac between CW9019 and RH4 cells. Log_2_ fold change was calculated from percent in peaks versus background.

